# Generalizable strategy to analyze domains in the context of parent protein architecture: A CheW case study

**DOI:** 10.1101/2022.05.07.491035

**Authors:** Luke R. Vass, Katie M. Branscum, Robert B. Bourret, Clay A. Foster

## Abstract

Domains are the three-dimensional building blocks of proteins. An individual domain can occur in a variety of domain architectures that perform unique functions and are subject to different evolutionary selective pressures. We describe an approach to evaluate the variability in amino acid sequences of a single domain across architectural contexts. The ability to distinguish different evolutionary outcomes of one protein domain can help determine whether existing knowledge about a specific domain will apply to an uncharacterized protein, lead to insights and hypotheses about function, and guide experimental priorities.

We developed and tested our approach on CheW-like domains (PF01584), which mediate protein/protein interactions and are difficult to compare experimentally. CheW-like domains occur in CheW scaffolding proteins, CheA kinases, and CheV proteins that regulate bacterial chemotaxis. We analyzed 16 domain architectures that included 94% of all CheW-like domains found in nature. We identified six *Classes* of CheW-like domains with presumed functional differences. CheV and most CheW proteins contained *Class* 1 domains, whereas some CheW proteins contained *Class* 6 (∼20%) or *Class* 2 (∼1%) domains instead. Most CheA proteins contained *Class* 3 domains. CheA proteins with multiple Hpt domains contained *Class* 4 domains. CheA proteins with two CheW-like domains contained one *Class* 3 and one *Class* 5.

We also created SimpLogo, an innovative method for visualizing amino acid composition across large sets of multiple sequence alignments of arbitrary length. SimpLogo offers substantial advantages over standard sequence logos for comparison and analysis of related protein sequences. The R package for SimpLogo is freely available.

## 1 INTRODUCTION

Many proteins consist of a chain of discrete domains. Protein domains are typically compact, independently folding units. Originally, protein domains were defined based on protease resistance; however, they are now recognized by characteristic amino acid sequence features. Individual domains possess structures and functions that are subject to natural selection and thus are discrete units of evolution. Domains were assembled in many permutations during evolution to create different proteins. Therefore, domains can be considered modular “building blocks” of proteins. The Protein Family (Pfam) database currently contains almost 20,000 different domains.^1^

Identifying domains is useful because once nature has evolved the solution to a problem (e.g., a protein domain that performs a specific function), the solution is frequently repeated. Thus, domain identity can provide useful general predictions about the structure and function of domains in experimentally uncharacterized proteins, as well as providing an effective means by which to compare proteins of known function. In recent years, more complex techniques have been proposed for comparing and quantifying homology between entire multi-domain proteins, including numerous alignment-free methodologies.^2,3^ Such approaches can successfully group functional proteins (such as kinases) at higher resolutions than standard sequence analyses by incorporating information on both catalytic domains and on attached accessory domains. However, much of the practical value of such predictions and our overall understanding of protein function still relies on the accuracy of discrete, individual domain descriptions (like those found in Pfam). Such domain families are often considered to be monolithic. While useful in some respects, in actuality substantial differences can be found between domains of the same type in different proteins. For example, response regulator proteins are formed by linking a receiver domain (Response_reg, PF00072) to a wide variety of output domains. At least three types of evidence demonstrate the differences between receiver domains in different types of response regulators: (i) phylogenetic analysis shows receiver domains co-evolve with their output domains;^4,5^ (ii) receiver domain:output domain interaction surfaces and receiver domain dimerization surfaces vary as a function of binding partner and whether a receiver domain exerts positive or negative regulation on its output domains;^6^ and (iii) receiver domains support phosphorylation and dephosphorylation reaction kinetics that differ by many orders of magnitude depending on the timescale of the associated biological function.^7–10^ Thus, receiver domains, though termed the same, differ substantially depending upon the domain architecture in which they are found.

Variation between different instances of a single specific protein domain raises multiple questions. If the structure and/or function of a domain are experimentally characterized in one protein, then how well does that information translate into useful insights about the same individual domain in another protein? In other words, how do protein domains themselves vary in the contexts of different domain architectures? Genome sequencing has far outstripped experimental capacity and provides a rich source of relevant information. Can sophisticated methods of amino acid sequence analysis identify contexts (i.e., location within domain architectures) in which current knowledge about a specific protein domain applies? Conversely, given the enormous scope of uncharacterized proteins and domain architectures, can we identify contexts where current knowledge about a specific domain is not applicable? Such a method would provide a way to inspire and focus future experimental investigations of protein domains in a manner that is most likely to be informative and may also lead to useful hypotheses and testable predictions. We describe here a generally applicable approach to analyze the variation in a specific protein domain as a function of the contexts in which the domain is found, revealing residue-level insights at a substantially higher resolution than a typical Pfam domain assignment.

As a test case to illustrate our method, we chose CheW-like domains (CheW, PF01584), which are part of the signal transduction pathways that regulate bacterial chemotaxis.^11^ CheW-like domains are known to act as scaffolds for various protein/protein interactions.^12,13^ The absences of an active site, substrate or ligand binding, or enzymatic activity make CheW-like domains challenging to study experimentally, because no simple assay exists by which to compare the properties of different CheW-like domains. One might envision measuring binding constants, but it would be difficult to untangle the contributions of the CheW-like domains and their partner proteins. Furthermore, CheW-like domains are used to assemble very large multi-protein complexes that couple chemoreceptors to kinases, so the biological relevance of the binding constant between an isolated CheW-like domain and its receptor or kinase partner is not immediately obvious. Finally, evolutionary constraints on CheW function are such that sequences can diverge substantially and still be functionally interchangeable.^14^ Thus, an objective method to classify CheW-like domains within different contexts would be particularly helpful to clarify the circumstances in which current knowledge is likely to be applicable and those which would benefit from further investigation.

CheW-like domains are found in three broad lineages of domain architectures: CheW scaffold proteins, which consist only of CheW-like domains; CheA kinases, which combine CheW-like domains with domains that provide kinase activity and phosphorylation sites; and CheV proteins, which combine CheW-like domains with receiver domains. Notably, the function(s) of CheV proteins are poorly understood^13,15^. A recent review of the diversity of bacterial chemotaxis systems asked, “Can efficient computational approaches be developed to distinguish … CheW from other CheW domain-containing proteins?”.^16^ Our results suggest that the answer is yes, and that not all CheW-like domains are alike.

## 2 MATERIALS AND METHODS

### 2.1 Protein sequence database

Protein sequences were initially obtained from the Pfam database (v.33).^1^ Any sequence found within the Representative Proteome 35 (RP35) to contain the CheW-like domain (PF01584) was extracted for further use, retaining only the CheW-like domain sequence.^17^ “35” refers to the co-membership threshold used to control the granularity of the sequence space represented. An RP set best represents the known sequence diversity of a group of similar proteomes/organisms. To construct the RP35 data set, available protein sequences for completed proteomes within the UniProtKB database were grouped based on their UniRef50 cluster membership. UniRef, or UniProt Reference Clusters, are sets of proteins grouped together by sequence in such a way as to produce tight clusters of orthologs/inparalogs, maximizing sequence space while minimizing redundancy; 50 refers to the sequence identity threshold used for the clustering procedure.^18^ We chose RP35 as the best compromise between information diversity and sequence space reduction. CheW-like domain sequences were grouped into *Architectures* based on the domain organization of the parent protein (Figure 2). Rare *Architectures* exhibited by fewer than 40 sequences were excluded from the analysis for practical reasons and because trends identified using such sparse data would likely be difficult to interpret with accuracy.

### 2.2 Sequence analysis and initial clustering

CheW-like domain sequences were collectively aligned to the accepted CheW-like domain HMM (PF01584) using HMMER3.^19^ The removal of poorly aligned sequences or subsequences has been shown to be beneficial for large-scale, downstream sequence analysis and phylogenetic profiling. To that end, we used trimAl (version 1.2) to remove spuriously aligned regions (using a column gap-threshold of 90%, assuming that the top 10% most “gappy” columns were mostly noise).^20^ Sequences with >50% gap-characters were also removed.

To identify the presence of the sequence subpopulations within our data set, we performed an initial cluster analysis on our full sequence set. Clustering of protein sequences can be performed in a number of ways, most commonly by calculating pairwise dissimilarity scores between sequences in an alignment and clustering the resulting matrix. Estimating protein sequence (dis)similarity requires the use of a substitution matrix to describe the rate or frequency at which a residue is substituted for another over time, simulating the process of evolution. We tested four commonly used evolutionary substitution models on our data set: GONNET, point accepted mutation 250 (PAM250; number of mutations per 100 amino acids), BLOcks SUbstitution Matrix 30 (BLOSUM30), and Jones-Taylor-Thornton (JTT).^21–24^ Each is tailored towards higher diversity sequence groups (the average sequence identity reported in the full Pfam alignment for CheW-like domain PF01584 is only ∼20%). We used the mat.dis() function from the bios2mds package in R (version 1.2.3) to calculate four pairwise-dissimilarity matrices from our full sequence alignment to use as inputs for *k*-medoids clustering.^25^

Clustering techniques like *k*-medoids requires an initial target number of clusters (the *k*-value) as input. An optimal *k*-value can be estimated by various metrics, such as the average silhouette width (a measure of how similar a point is to its own cluster compared to other clusters), but choosing the “correct” number of clusters is challenging, subjective and dependent on the overall goals of the analysis.^26^ *A priori*, we chose *k* = 21 (the total initial number of distinct architectural *Contexts*, including multiple domains in the same protein, used to partition the CheW-like domain sequences). This was an intentional overestimation of the true number of subpopulations. An optimal *k* value (∼3-4) was also estimated *a posteriori* using the average silhouette width method implemented in the pamk() function from the fpc package in R (version 2.2-9; *k*=3-10).^27^ Initial *k*-medoids clustering was performed using the pam() function from the cluster package within R (version 2.1.2; *k-*medoid clustering is sometimes called <u>P</u>artitioning <u>A</u>round <u>M</u>edoids).^28^

For convenience, we only show solutions derived from the GONNET substitution matrix in the main text. Solutions derived from the other three models are featured in the Supporting Information. The similarity of the partitions generated using the four substitution models was quantified using both the Adjusted Rand Index (ARI)^29,30^ and Normalized Mutual Information (NMI),^31,32^ commonly used metrics when comparing clustering schemes. Each of the four solutions was highly similar, with ARI scores ranging from 0.6-0.7 and NMI scores of 0.7-0.8, suggesting a relatively stable and generalizable underlying data structure.

### 2.3 Multidimensional scaling analysis

The pairwise dissimilarities between the medoids for each of the 21 identified *Clusters* were extracted and used as input for analysis with multidimensional scaling (MDS). MDS is a multivariate dimensionality reduction and visualization technique that takes pairwise similarities/dissimilarities/distances for a data set and attempts to reconstruct a simplified map that maintains the relative positions of the data points and is easily interpretable (i.e., in a lower *k-*dimensional space). Two main types of MDS exist: metric MDS, which focuses on the “distances” in the matrix being analyzed, and non-metric MDS, which assumes that onl the relative rankings of the distances are known.

Due to its simplicity and previous success when applied to protein sequences, we chose metric MDS^25^ to visualize the results of the initial *k*-medoid clustering procedure. For the subsequent subsampling steps (described below), we first tested an implementation of the metric MDS algorithm featured in the wcmdscale() function from the vegan package in R (version 2.5-7).^33^ However, other forms of MDS, such as the non-metric approach, may offer additional advantages under the appropriate circumstances. Examination of our MDS solutions indicated that a significant proportion of the total eigenvalues were negative (∼15.4-15.7%). Though the individual negative eigenvalues were miniscule compared to the positive entries, it was possible that our initial dissimilarities were non-Euclidean in nature, and non-metric MDS might offer additional insight. Consequently, we tested a non-metric implementation of the MDS algorithm using the metaMDS() function, also from the vegan package (using parameters: *trymax = 2000, noshare = F, maxit = 9999, sratmax = 0.99999999, k = 3, model = “local”, autotransform = F, wascores = F, center = T, pc = T*) ^33^. The resulting MDS plots (included in the Supporting Information) were visualized using the ggplot2 package in R (version 3.3.3).^34^ While we encountered significant challenges achieving convergence (likely due to the still relatively large sequence set), the non-metric MDS solutions were largely reminiscent of the basic layouts generated using the metric algorithm, even when sampling smaller subsets of sequences.

### 2.4 Sequence subsampling strategy

A primary technical disadvantage of the MDS technique is its substantial computational cost. The time complexity of standard metric MDS is proportional to the square of the input size, indicating significant temporal and technical requirements when analyzing a high-dimensional data set with thousands of data points.^35^ Thus, while we could effectively apply MDS to the cluster medoids from our limited initial analysis, the approach becomes impractical when analyzing a larger, more complete data set like our full dissimilarity matrix featuring ∼8500 CheW-like domains.

One popular approach to dealing with large, unwieldy data is random subsampling, called stratified sampling when working with distinct groups or strata within the original population. However, for this strategy to be effective, adequate representation of each stratum is essential. Guided by our initial *k*-medoids clustering results (which effectively revealed any strata within the sequence data), each architectural *Context* was treated as a separate homogeneous class, except for CheW.I, which was split into CheW.IA, CheW.IB and CheW.IC *Types*. CheW-like domains from parent *Architectures* featuring multiple domains were split into separate *Contexts* with appropriate suffixes added relative to their N- to C-terminal position in the parent protein. Two main approaches to stratified subsampling exist: randomly selecting a predefined number of observations from each defined subgroup; or randomly selecting a number of observations proportional to the original subgroup sizes. Given the wide disparity in the number of CheW-like domain sequences of each *Type*, we chose the former to facilitate computational efficiency. The mat.dis() function from the bios2mds R package was again utilized for the iterative sampling procedure.^25^ Each subsampling round randomly selected 25 sequences from each of the 23 *Types* (without replacement; smaller subsets were also explored with similar results). Sampling was repeated 10 times (575 distinct sequences per iteration), generating a total of 10 replicate sequence sets for each of the four previously described substitution models, to assess the stability of the various randomized MDS solutions. We also included the CheW-like domains of 18 well-studied reference proteins as an internal constant for each iteration (data not shown). Pairwise dissimilarity matrices were estimated for each replicate set for further use during the MDS analysis described above.

### 2.5 Conservation analysis and SimpLogo creation

All CheW-like domain sequences were first aligned collectively to the conserved CheW-like Hidden Markov Model (HMM) profile (PF01584) obtained from the Pfam database. Separate sequence alignments were then segmented based on corresponding *Types* of CheW-like domains. For each alignment, the *Escherichia coli* CheW sequence was inserted at the top for use as a reference. Columns introducing gaps (relative to the *E. coli* CheW sequence) were removed. Position-wise count matrices and corresponding frequencies were generated for each alignment using the Biostrings and motifStack packages in R.^36,37^ Physicochemically similar residue types were combined to reduce visual complexity of the final representation, creating nine amino acid groups (including gap characters): positively-charged side chains (His/Lys/Arg), negatively-charged side chains (Asp/Glu), small volume side chains (Ala/Gly), polar side chains (Gln/Asn/Ser/Thr), large hydrophobic side chains (Met/Val/Leu/Ile), gaps (-), aromatic side chains (Phe/Tyr/Trp), Pro, and Cys. Each amino acid category was assigned an “idealized” color and converted to tristimulus values in the CIE XYZ color space. XYZ space is linear and non-negative (as opposed to traditional RGB space), making it simple to mix colors. Final values were calculated by weighting (multiplying) the idealized colors of each amino acid group by their relative frequencies and summing the results for each position. Logos were visualized using geom_tile objects (from the ggplot2 suite), with colors and heights for each position based on the aforementioned blended tristimulus values and their overall gap character frequency, respectively. “Idealized” (i.e., color at 100% frequency) color strips for the top two most frequent residue types were added to each position (at the top and bottom of each tile) to mitigate uncertainty between true qualitative colors and blended hues. Position-wise information content (in bits) was estimated for each alignment again using motifStack and visualized with a line trace at the top.^37^ To create traditional sequence logos (with similar dimensions, for comparison purposes), the Logolas R package was used with default settings.^38^ An R package for SimpLogo was created and is freely available (along with an example and the sequence alignments used to generate Figure 5) in the Github repository https://www.github.com/clayfos/SimpLogo (v.0.1.1, https://doi.org/10.5281/zenodo.6323714).

### 2.6 Protein homology modeling and structural analysis

Crystal structures of the CheW-like domains from experimentally well-studied *E.* coli CheW or CheA proteins are not available. To visualize the relevant functional surfaces of the reference *E. coli* CheW-like domains CheA-P5 and full CheW protein, we modeled trimeric CheW:CheA-P5:receptor complexes (capturing both interface 1 and 2) based on the crystal structure of the *Thermotoga maritima* multimer featuring CheW, CheA-P5 fragment, and the chemoreceptor TM14 (PDB ID 4JPB).^39^ We first generated homology models of *E. coli* CheA-P5 (residues 514-640; using the crystal structure of *T. maritima* CheA PDB ID 1B3Q^40^ as a template) and CheW (residues 18-154; using the solution structure of *E. coli* CheW PDB ID 2HO9 as a template) with the Phyre2 web portal.^41^ We then superimposed the top models against the corresponding *T. maritima* molecules in the crystal structure (PDB ID 4JPB^39^) using PyMOL (v.2.3.3).^42^

## 3 RESULTS

### 3.1 CheW-like domains are incorporated into dozens of distinct domain *Architectures* in nature

We surveyed the known domain architectures into which CheW-like domains are incorporated, which provided initial sequence groupings for further investigation. We extracted the sequences of all proteins containing a CheW-like domain (PF01584; 8484 sequences containing 9094 unique CheW-like domains) from the Pfam database (v.33)^1^ using the Representative Proteome 35 (RP35) sequence set.^17^ The use of RP35 provided an efficient and representative data set while substantially reducing the sequence space and complexity required for the analysis. CheW-like domain sequences from rare architectures (<40 member sequences) were excluded because we anticipated that their interpretation would be too speculative. The remaining sample contained 8581 CheW-like domains (∼94% of original) in 8061 proteins (∼95% of original). Additional curation of sequences described in Methods (e.g., for excessive gap character) resulted in final analysis of 8528 sequences from CheW-like domains.

We employ the capitalized and italicized terms *Architecture*, *Context*, *Cluster*, *Group*, *Type*, and *Class* to refer to the specific hierarchy used to categorize sequences of CheW-like domains in this study. The relationships between the classification terms, as well as the flow of our sorting scheme, are outlined in Figure 1. We first categorized CheW-like domain sequences into three “lineages” based on the architecture and putative role(s) of their parent protein (Figure 2). Our nomenclature for *Architectures* containing CheW-like domains used the accepted lineage of the parent protein (e.g., CheA) as a prefix and a Roman numeral as a suffix. CheA-lineage *Architectures* were numbered by decreasing membership size, starting with Roman numeral I. Additionally, *Architectures* containing two or more CheW-like domains (e.g., CheA.V from Figure 2) were split into their constituent CheW-like domains and analyzed as separate sequence *Contexts*. An additional suffix of an Arabic numeral identified the “position” of the CheW-like domain relative to the N-terminus and the other CheW-like domains in the parent protein, e.g., CheA.V.1 (*Contexts* are not labeled in Figure 2). The CheA-lineage was the most diverse, with a dozen major *Architectures*. In addition to CheW-like domains, most CheA *Architectures* also contained HATPase_c domains to bind ATP and catalyze phosphorylation, and histidine phosphotransferase (Hpt) domains to provide a phosphorylation site.

**FIGURE 1.**
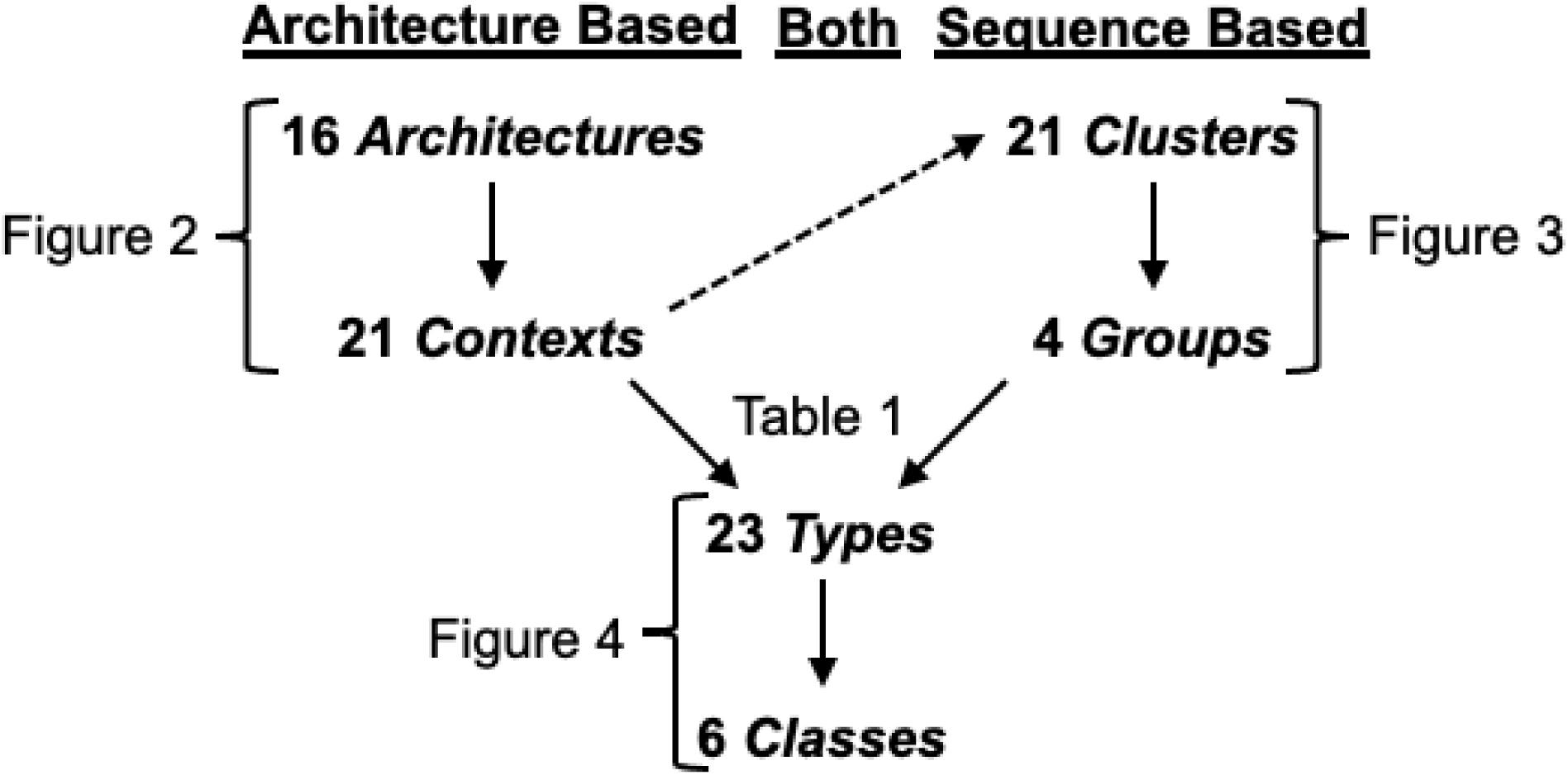
Hierarchy of classification terms and sorting workflow. The initial steps were derived purely from domain *Architectures*, which may contain CheW-like domains in more than one *Context*. The number of *Contexts* defined (dashed arrow) the number of *Clusters* into which sequences of CheW-like domains were initially sorted. However, sequences in a given *Context* sorted into different *Clusters* and vice versa. The *Clusters* sorted into larger *Groups* of sequences. Comparison of architectural *Contexts* with sequence *Groups* revealed that all but one *Context* included *Clusters* within a single *Group*. One *Context* contained *Clusters* that spanned three *Groups* and so was split into three *Types*, whereas all other *Contexts* became a single *Type*. Minor *Clusters* of sequences associated with each *Context* were discarded from further analysis during the transition to *Types*. The *Types* sorted into larger *Classes* of sequences associated with one or more architectural *Contexts*. Figure 5 illustrates the final *Types* and *Classes* but was not part of the classification process. See text and indicated Figures or Table for details.

**FIGURE 2.**
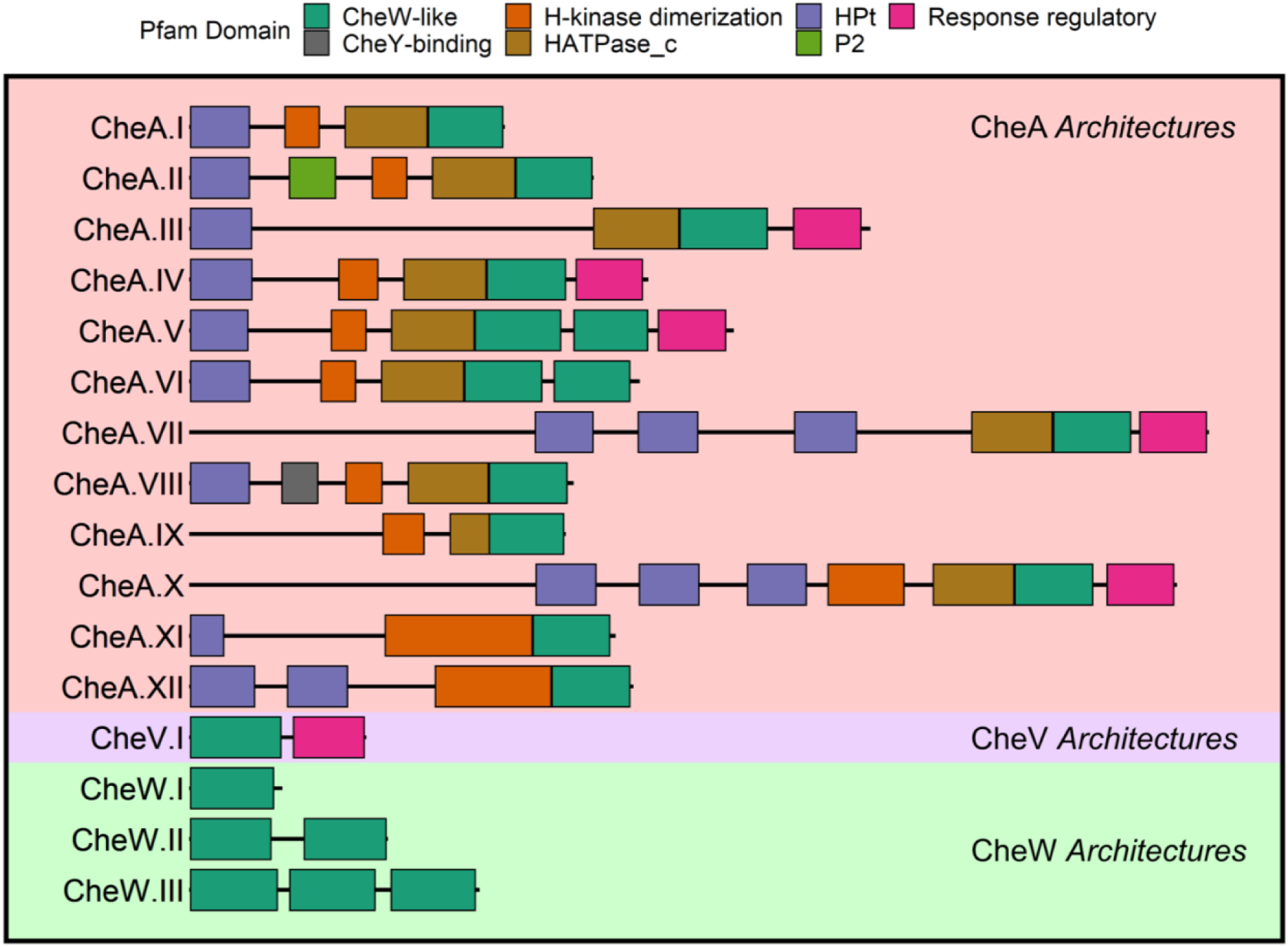
Schematic of the most abundant CheW-containing domain *Architectures* seen in nature. Sixteen distinct architectural variants containing the CheW-like domain were identified within the Pfam database (≥40 sequences each, ordered by decreasing abundance within each shaded group). Sequences for their corresponding CheW-like domains were extracted and initially classified according to the schematic. For parent proteins containing multiple CheW-like domains, CheW-like domains were initially classified according to the position (*Context*) of the domain, resulting in one sequence entry per CheW-like domain. Representative diagrams are shown for each *Architecture* (from left to right, N- to C-terminal). Pfam domain IDs included are as follows: CheW-like (PF01584); H-kinase dimerization (PF02895); Hpt (PF01627); Response regulatory (PF00072); CheY-binding (PF09078); HATPase_c (PF02518); P2 (PF07194). Individual schematics are not necessarily drawn to scale.

Although several CheV-lineage architectures (named for the prototypical *Bacillus subtilis* CheV protein^43^) were detected in the RP35 set, the only organization incorporating an appreciable number of sequences (>40) was the prototypical form, comprised of a N-terminal CheW-like domain and a C-terminal receiver domain, which was termed the CheV.I *Architecture* for consistency. The CheV-lineage was the least abundant of the three architectural lineages.

CheW-lineage architectural variants constituted the largest collection of sequences, with over half of the entire data set attributable to prototypical CheW proteins comprised of a single CheW-like domain, from which the *Architecture* gets its name (CheW.I). Additional rare (∼1 to 3% the number of CheW.I) *Architectures* featured two or three CheW-like domains fused together (termed CheW.II and CheW.III respectively) with no other detectable domains present. Individual CheW-like domains of CheW.II and CheW.III *Architectures* were separated into distinct *Contexts* for further analysis, doubling or tripling the number of sequences for the corresponding *Architecture* (i.e., CheW.II.1, for the first position CheW-like domain, CheW.II.2 for the second position CheW-like domain, etc.).

Although we focused only on the more abundantly represented architectural variants, CheW-like domains are ubiquitous and can be found embedded in dozens of distinct *Architectures* encompassing nearly every component and layer of the microbial chemotaxis system.^44^ We briefly investigated the sequence groups that failed to pass our chosen thresholds. The majority of these “rare” *Architectures* fell within the CheA-lineage (containing anywhere from zero to 10 Hpt domains). Atypical, full-length CheA-lineage proteins were also present, fused to CheB, CheC, CheX, or MCP domains. In addition to a CheW.IV *Architecture*, CheW-like domains were detected fused to CheC, CheR, CheX, CheZ, Hpt, or receiver domains, as well as PAS domains that might provide input information to chemosensory systems. Finally, CheW-like domains were infrequently found in proteins with a variety of other domains that have no obvious connection to chemotaxis.

### 3.2 CheW-like domains segregated into distinct sequence *Groups* as a function of parent domain *Architecture*

We expected that CheW-like domains extracted from a common architectural *Context* would more closely resemble each other rather than the same domains obtained from contrasting architectural variants. We based our assumption on existing knowledge of microbial response regulator receiver domains, which cluster phylogenetically predicated on the presence and/or type of an attached output domain.^4,5^ However, the full dataset contained >8500 CheW-like domain sequences, making analysis and interpretation challenging. Ideally, each CheW-like domain *Context* would behave as a homogeneous sequence set, and a basic stratified sampling strategy could be used to select a more representative and manageable population for study. While the assumption of homogeneity seemed reasonable (shared architecture typically implies common function), we wished to first gain insight into the underlying structure of the sequence space by performing unsupervised clustering on the full list of sequences. This allowed us to more accurately assess and incorporate the heterogeneity of the data into our approach. Essentially, our overall approach incorporated a simplified multistage sampling scheme, first clustering the entire population to reveal major strata within each architectural *Context*, then randomly sampling from within these strata, ensuring that all major sequence groups were represented. We chose to use a *k*-medoids approach for the initial clustering procedure to mitigate the difficulty of visualizing >8500 data points. The *k*-medoids algorithm is a classical clustering technique that partitions the data into *k* number of *Clusters*, attempting to minimize the distance between data points and the center of their relevant *Cluster*. Each *Cluster* is represented by a single central data point called a cluster medoid, which allowed us to visualize the *Clusters* using just the medoid sequences, rather than the entire data set.

A representative multi-dimensional scaling (MDS) plot describing just the 21 extracted medoids (based on the dissimilarities calculated using the GONNET model) is shown in Figure 3. Additional plots generated using dissimilarities calculated with three other substitution models are included in Figure S1 (all four partitions were highly similar). Letters naming the *Clusters* for each solution were assigned based on decreasing number of sequences in the *Cluster*. There were three substantial *Groups* of *Clusters* of CheW-like domains (labeled 1, 3, or 4 in Figure 3), and another smaller, intermediate *Group* separating *Groups* 1 and 3 (labeled 2 in Figure 3, more apparent when considering only the first dimension).

**FIGURE 3.**
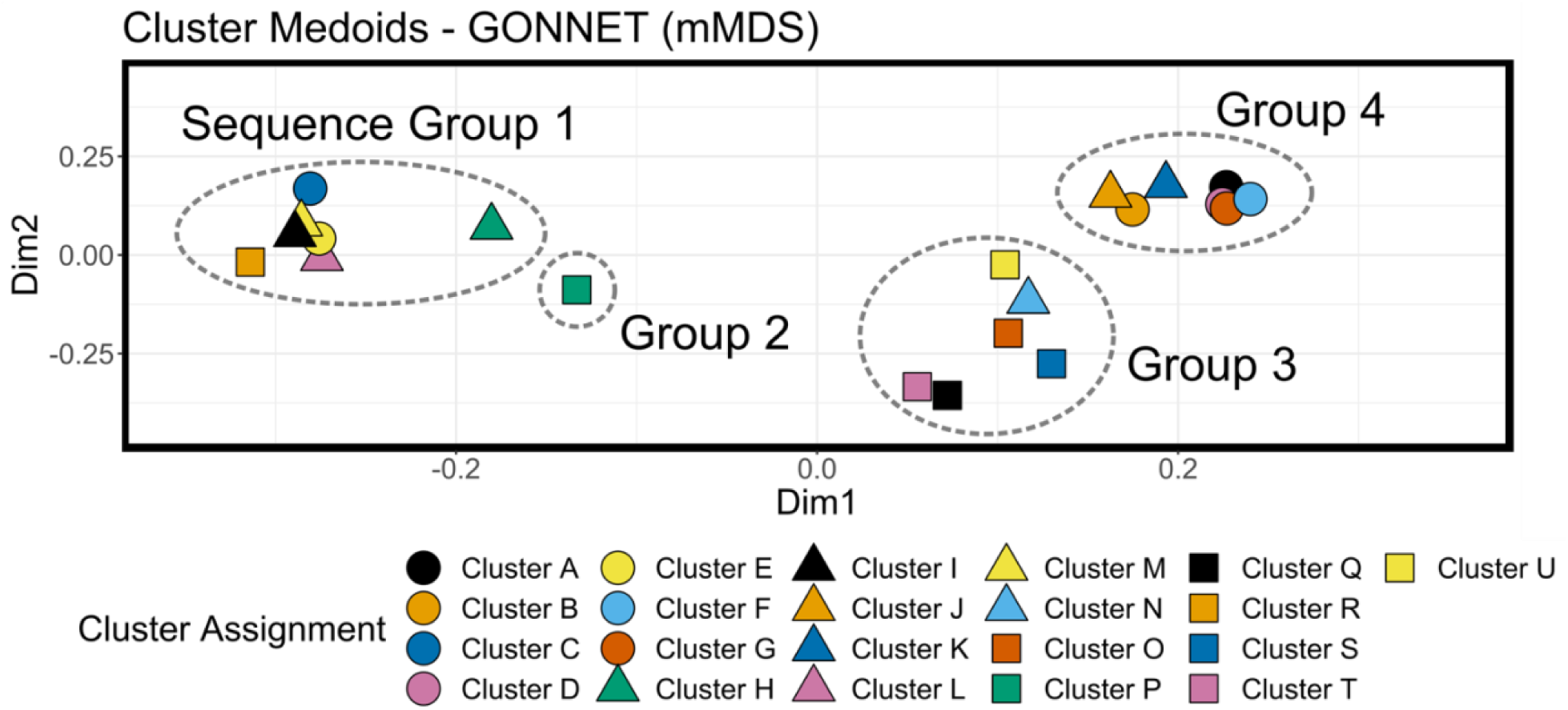
Cluster analysis and multidimensional scaling of representative sequences of CheW-like domains reveals *Groups*. Multidimensional scaling plots showing representative medoids from each of the 21 *Clusters*, generated using dissimilarity matrix calculated with the GONNET substitution model. Each clustering attempt reconstituted roughly the same data structure, consisting of three main *Groups* (numbered 1, 3 and 4) of *Clusters* along with a weaker, more intermediate *Group* (numbered 2) intersecting *Groups* 1 and 3. While the colors and specific *Cluster* designations may shift between analyses (overall *Cluster* sizes were slightly altered depending on the substitution matrix used, affecting *Cluster* names), the actual partitioning of CheW sequences was highly consistent, regardless of the substitution model used (see Figure S1).

We next examined the composition of the individual *Clusters* to determine if the original architectural *Contexts* of CheW-like domains separated into multiple stable strata. Although each medoid was representative of the dissimilarity data, a *Cluster* may contain multiple domain *Contexts*. Understanding the composition of each *Cluster* would allow us to properly organize sequences for analysis on the more complete set, rather than on just the representative medoids, ensuring that no distinct *Contexts* would be underrepresented. Table 1 assigns *Clusters* to the four main *Groups* indicated in Figure 3 and contains a detailed breakdown of the architectural *Contexts* of the sequences in each *Cluster*.

**Table 1.**
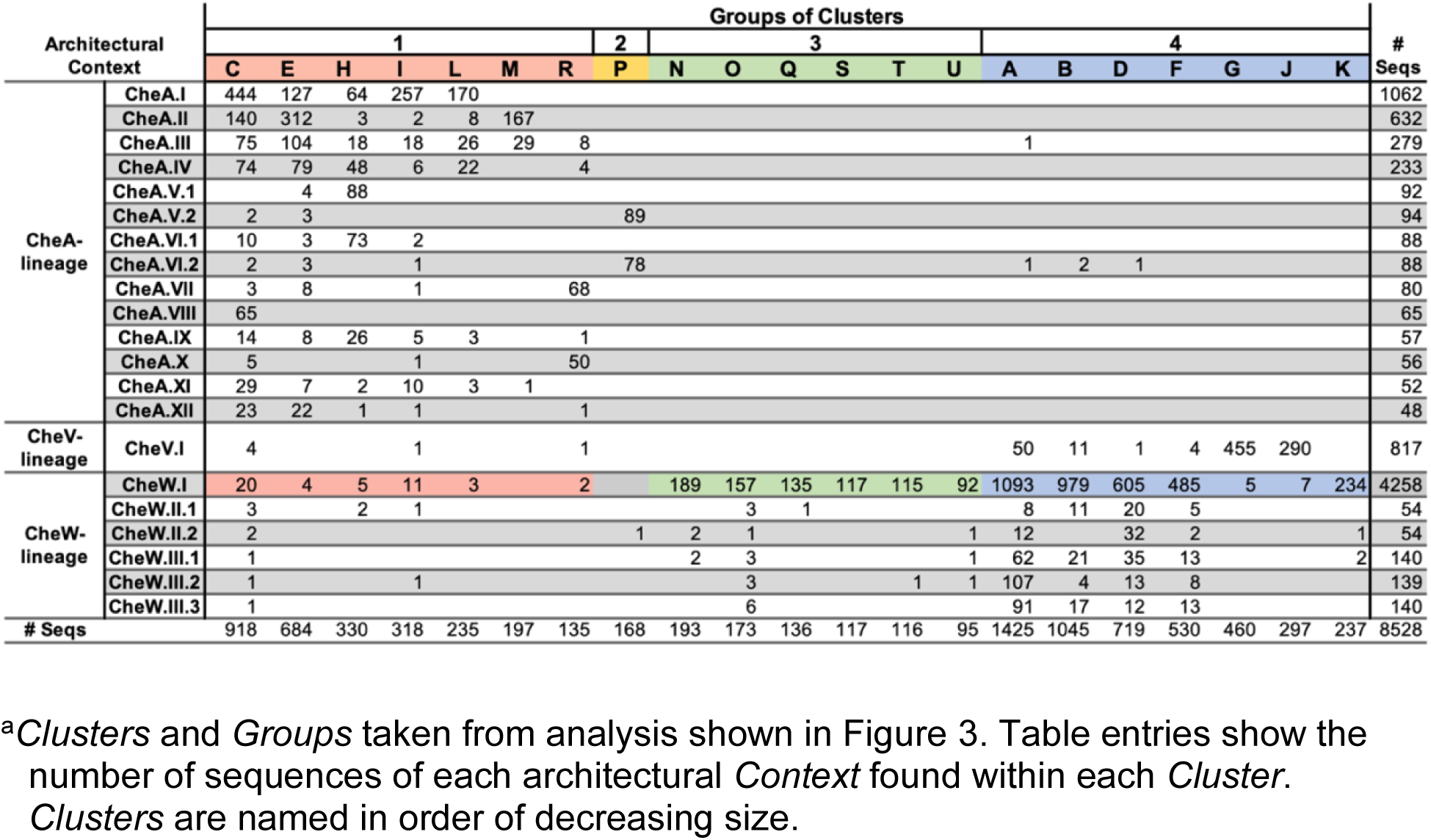
Distribution of CheW-like domain *Context*s across *k*-medoid *Clusters*^a^.

Multiple patterns are apparent in Table 1. First, sequences form *Clusters* and *Clusters* form *Groups* primarily based on parent protein lineage. Most or all sequences within a *Cluster* are from either CheA proteins or CheW/CheV proteins, not both. Most CheW-like domains from CheA proteins are in *Group* 1 and most CheW-like domains from CheW and CheV proteins are in *Group* 4. Closer inspection reveals additional structure in the data:

- The “second” position CheW-like domains from *Architectures* CheA.V and CheA.VI formed their own distinct sequence *Group* 2, made up of *Cluster* P.
- CheW-like domains from *Architectures* CheA.VII and CheA.X comprised the bulk of *Cluster* R, the smallest *Cluster* in *Group* 1.
- About 80% of domains from CheW.I *Architectures* and greater than 90% of domains from CheW.II and CheW.III *Architectures* were in *Group* 4.
- About 20% of domains from CheW.I *Architectures* and 5% of domains from CheW.II and CheW.III *Architectures* were in *Group* 3, possibly distinct from the majority of CheW-lineage domains.
- About 1% of domains from CheW.I *Architectures* and 2% of domains from CheW.II and CheW.III *Architectures* were in *Group* 1, suggesting similarity to the CheA-lineage.
- About 99% of CheW-like domains from CheV.I *Architectures* were in *Group* 4, suggesting that the presence of the attached receiver domain has little effect on the primary sequence of the CheW-like domain in CheV proteins.
- Similar to CheW proteins, about 1% of CheW-like domains in CheV.1 *Architectures* were in *Group* 1, suggesting similarity to the CheA-lineage.

Based on the analysis of Table 1, we assumed for further processing that each architectural *Context* could be reasonably treated as a single homogeneous entity, except for CheW.I. We split CheW.I into three distinct *Types* designated CheW.IA (sequence *Group* 3), CheW.IB (sequence *Group* 4), and CheW.IC (sequence *Group* 1). Remaining outlier sequences in each architectural *Context* (typically a small number of sequences in each *Context* assigned to substantially different *Clusters* than the majority) were excluded from the analysis as noise. We believe that our evaluation produced an accurate summary of the underlying data structure, though we acknowledge the somewhat subjective nature of cluster analysis and that alternative perspectives may offer additional valid insights. To explore the data at higher resolution and understand sequence variation within the *Clusters*, we next performed MDS analysis on a larger, randomly sampled subset of our sequence data, now organized into 23 *Types* (due to CheW.I sub-designations) instead of 21 *Contexts*.

### 3.3 Random stratified subsampling facilitates efficient multi-dimensional scaling analysis on large protein sequence sets and revealed *Classes* of CheW-like domains

Our initial cluster analysis (Figure 3 and Table 1) revealed how to organize sequences of CheW-like domains into 23 *Types* that should ensure all major subpopulations receive appropriate representation in a random stratified subsampling scheme described in Methods. Figure 4 shows the top three dimensions of one representative solution derived from the GONNET substitution matrix. The remaining nine iterations, as well as solutions calculated using randomly sampled subsets for the other three substitution matrices described in Methods, are in Figures S2-S13. The solution shown in Figure 4 revealed similar relationships to those found in Figure 3/Table 1, which we now designate as *Classes* (sub-sampling MDS) rather than *Groups* (*k*-medoid MDS). On the right area of the plot (Left and Center Panels), most CheW-lineage and CheV-lineage sequence *Types* (*Class* 1) clustered in a densely packed region. A large agglomeration of sequences from CheA-lineage *Types* (*Class* 3) was near the left side of the plot area (Left and Center panels). The CheA.V.2 and CheA.VI.2 *Types* (*Class* 5) were markedly different from the other CheA-lineage *Types*, consistent with previous assignment to *Cluster* P as the sole member of *Group* 2 in Figure 3/Table 1. Additional differentiation within the CheA-derived *Types* could be observed more effectively in the Center and Right Panels (incorporating Dimension 3), with the CheA.VII and CheA.X *Types* (*Class* 4) separating from the main body. This supports the clustering results presented in Table 1 and Figure 3, with the majority of CheA.VII and CheA.X sequence *Types* found in the somewhat independent *Cluster* R. Sequences belonging to the rare CheW.IC *Type* (*Class* 2) were spread across Figure 4, alongside *Class* 1 CheW/CheV sequences, *Class* 3 CheA sequences, and in between (Left and Center Panels). In Figure 3/Table 1, a large subset of CheW-lineage sequences (*Group* 3) separated from most CheW- and CheV-lineage sequences (*Group* 4). The *Group* 3 sequences are termed *Type* CheW.IA (*Class* 6) in the analysis of Figure 4 and surprisingly did not separate from the main CheW/CheV *Types* (*Class* 1).

**FIGURE 4.**
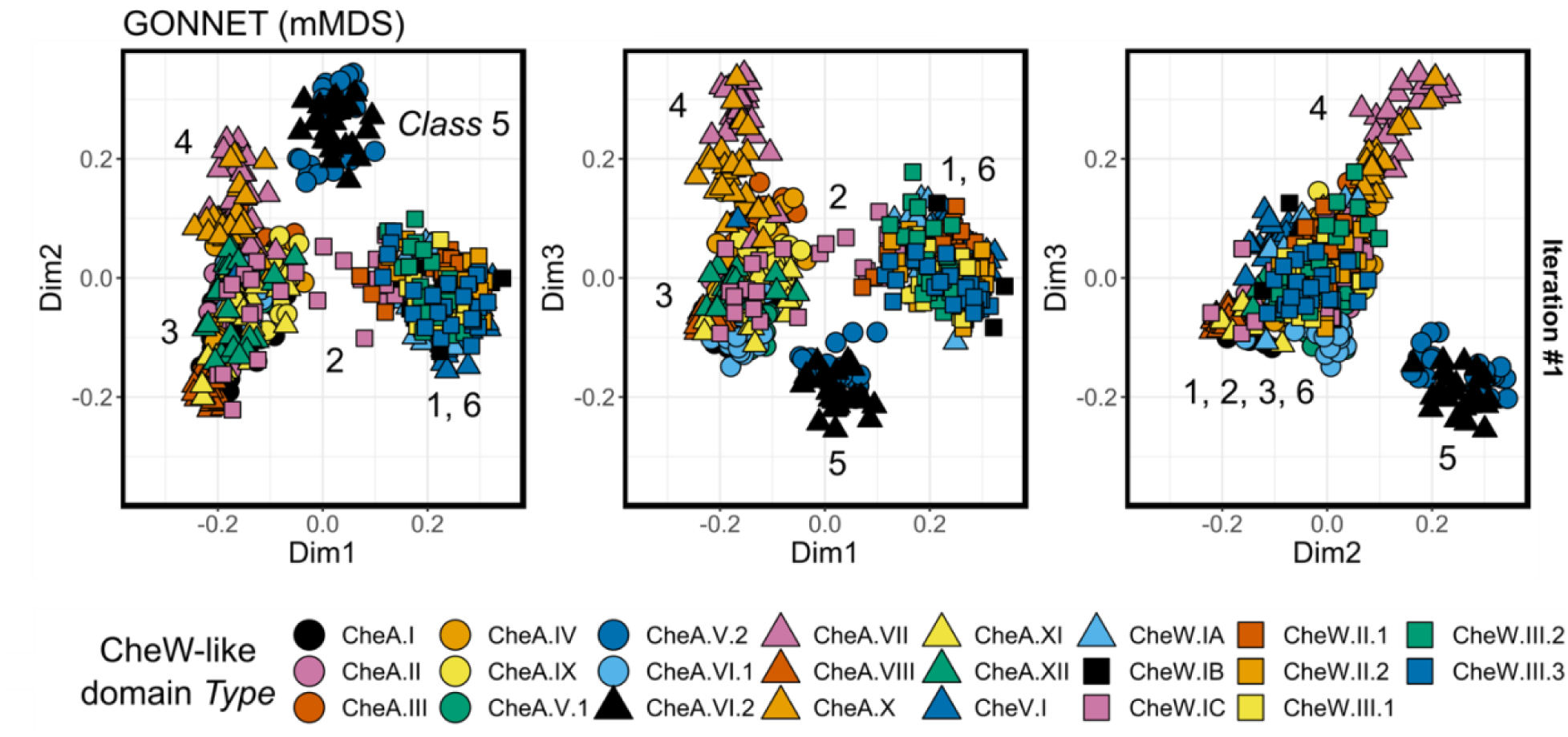
Metric multidimensional scaling plots showing 25 randomly subsampled sequences from each of the 23 CheW-like domain *Types*. The top three dimensions (Dim1, Dim2, Dim3) were included for visualization purposes. Analysis was performed on dissimilarities calculated using the GONNET substitution model. The location of *Types* belonging to each of six *Classes* are indicated. Additional sampling iterations are shown in Figures S2-S4. Sampling iterations with other substitution models are shown in Figures S5-S13.

We also tried non-metric MDS as described in Methods and shown in Figures S14-S25. We concluded that the data structures were likely consistent irrespective of the MDS algorithm. Given the clear sequence heterogeneity within the CheW-like domains, we next sought to determine factors within the domain itself that distinguished the various *Types*.

### 3.4 SimpLogo: A simplified method for visualizing large numbers of distinct protein sequences aligned over many positions

Arguably the most popular method for comparing sequence groups and their alignments is the sequence logo. In this context, a logo is a graphical representation depicting sequence conservation by stacking relevant characters, such as single-letter codes for amino acids or nucleotides, at each position in an alignment.^45^ The relative heights of the characters are proportional to their frequency at the given position. Additionally, the overall height of a stack is often scaled based on the information content stored at that position within the alignment. Typical applications for amino acid sequence logos include visualizing functional motifs or binding sites within a protein family/group. However, the volume of information and the graphical complexity of a logo often makes it challenging to interpret when examining regions longer than two dozen positions. Sequence logos are less practical when comparing a large group of separate alignments (>12) with many positions (>50). A more ideal and complementary visualization technique would allow us to first evaluate the macroscopic characteristics of each sequence group. Subtle differences could then be examined in greater detail for specific regions using a more traditional approach like a true sequence logo, or even the original sequence alignment. Here, we describe a visualization format (colloquially referred to as SimpLogo) designed to compare the CheW-like domain sequences from the numerous *Types* described above. We sought to reduce the 23 individual sequence alignments (*Types*) into simplified, pseudo-one-dimensional representations. We accomplished this by weighting and blending qualitative colors assigned to specific amino acid residue types (grouped by physicochemical properties into eight classes) based on their frequency at a given position within the alignments (Figure 5). We also modified the overall height based on its proportion of gap-character (details in Methods). This produced a single row of colored “boxes” for each sequence *Type*, reducing a complex problem to basic pattern recognition. To differentiate between secondary colors and a near-equal mixing of primary colors, colored strips were included on the top and bottom of each box, indicating first and second most frequent residue types. The simplified scheme allowed for the simultaneous comparison of dozens of distinct sequence families with lengths over many positions (>100) and encoding substantially more information about intermediate frequencies than simple consensus sequences. Additionally, we created traditional sequence logos (using the Logolas package in R) for the same alignments using similar image dimensions for comparison (see Figure S26). Highly conserved/informative positions are identifiable in the traditional logo, but the overall lack of space and reduced character sizes often make interpretation of specific positions challenging, especially for the lower frequency/less conserved positions with smaller stack heights. The patterns of gap characters (indicated by dashes) are also more challenging to interpret when compared to the SimpLogo representation in Figure 5. One could imagine how difficult interpretation would become with double the number of sequence groups or sequence lengths using traditional methods. Therefore, we envision the new SimpLogo visualization technique being used not as a replacement for traditional sequence logos, but as a supplementary technique for large scale exploratory analyses.

**FIGURE 5.**
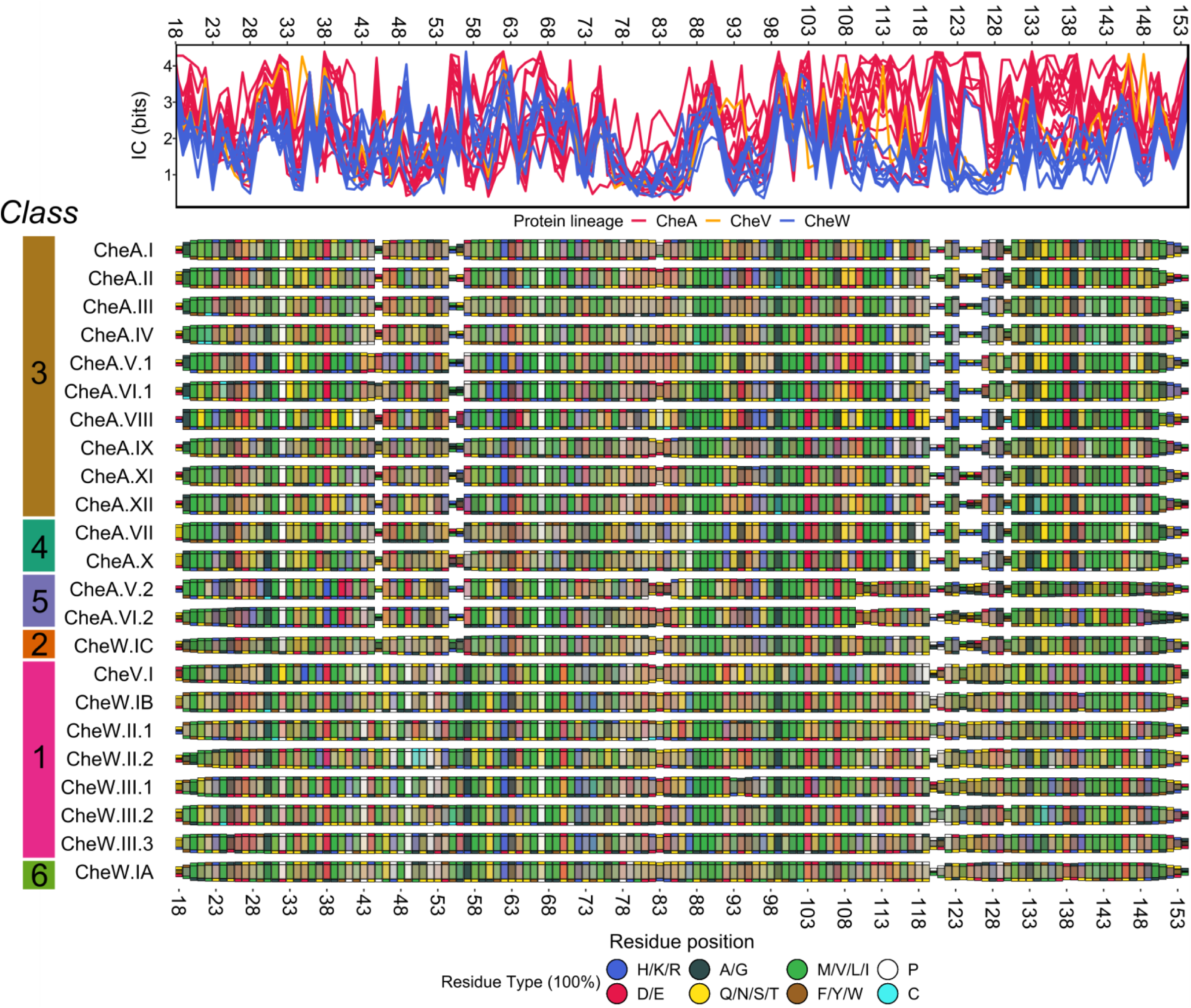
SimpLogo representation of major CheW-like domain *Types*. Rows correspond to domain *Types*. Sequences were aligned to the CheW-like domain HMM (Pfam ID PF01584) and separated by *Type*. Residues with physicochemical similarities were merged and assigned colors. Idealized colors (legend) were weighted by residue frequency and mixed to produce a colored box at each position. Heights were scaled by gap percentage. To differentiate between secondary colors and a near-equal mixing of primary colors, colored strips were included on the top and bottom of each box, indicating first and second most frequent residue types respectively. *Types* were organized by *Class* in the figure. Position-wise information content (in bits) was calculated for each alignment (top). Numbering is for *E. coli* CheW, framing our observations with reference to the well-studied prototypical protein.

### 3.5 *Classes* of CheW-like domains display characteristic gap patterns and residue identities

The SimpLogo representation in Figure 5 revealed multiple trends. Overall, the relationships displayed in Figure 5 mimicked the trends seen in Figure 4, with most of the CheA-lineage *Types* clustered tightly (*Class* 3), the exceptions being the CheA.VII/CheA.X (*Class* 4) and CheA.V.2/CheA.VI.2 (*Class* 5) *Types*. Most CheW-lineage *Types* (*Classes* 1 and 6) and the CheV-lineage *Type* (*Class* 1) appeared similar, except for the CheW.IC *Type* (*Class* 2), which was intermediate in character between CheA and CheW *Types*.

Figure 5 is information dense, so interpretation is facilitated by provision of appropriate context. *E.* coli CheW and CheA proteins are experimentally well-studied, but crystal structures of the corresponding CheW-like domains are not available. To provide a structural frame of reference for our analysis, we constructed simple homology models of *E. coli* CheW and the CheW-like domain (termed P5) of CheA based on experimentally determined crystal structures of the corresponding *T. maritima* domains as described in Methods. For convenient cross-referencing, Table S1 contains a detailed list of putative functions for relevant residues of CheW-like domains based on various literature sources and our homology models. We assumed, for the sake of speculation, that the analogous interaction surfaces and behaviors are largely maintained throughout the various CheW-like domain topologies found in nature and represented in Figure 5. A glance reveals that the gap characters (box heights) of the CheA-, CheV-, and CheW-lineages were different, which likely played an important role in differentiating the sequence *Types*, particularly at positions 45-56 and 120-131. Similarly, there were clear differences in residue identities, especially involving *Class* 5 domains. In Text S1, we describe observations concerning gap patterns and residue identities at length, and also suggest possible functional implications. These descriptions illustrate the detailed information and hypotheses that can be derived from analysis of SimpLogos in conjunction with existing experimental data. However, the primary purpose of this report is to describe our generally applicable strategy to analyze protein domains in different architectural contexts and, as summarized in the Discussion, the higher order conclusions that can be drawn from sorting a single Pfam domain into different *Classes*.

## 4 DISCUSSION

### 4.1 Implications of six CheW-like domain *Classes*

The existence of *Class* 1, which encompasses CheV and most CheW proteins, has multiple implications. First, evidence exists that CheV proteins can function as scaffold proteins similarly to CheW proteins^12,46–48^; however, more detailed information about the functions and mechanisms of CheV is sparse.^13,15^ It is tempting to extrapolate what is known about response regulator proteins (composed of N-terminal receiver and C-terminal output domains) to formulate hypotheses about CheV (composed of C-terminal receiver and N-terminal CheW-like domains). However, our observation that CheW-like domains in CheV proteins were similar to those in CheW proteins constrains such hypotheses by suggesting that any interactions between the receiver and CheW-like domains in CheV proteins are not sufficiently specific to be subject to selective pressure for evolutionary optimization. There are at least three categories of hypotheses about CheV inspired by response regulators:

- The receiver domain might alter the conformation of or occlude access to the CheW-like domain, depending on the phosphorylation state of the receiver. Alexander *et al.* suggested that such phosphorylation-mediated changes could explain the experimentally established role of CheV phosphorylation^49,50^ in adaptation by changing the conformation of the CheW-like domain to alter the degree of coupling between receptor and kinase function.^15^ Our findings make this otherwise reasonable mechanism now seem unlikely. In *Bacillus subtilis* CheV, the CheW-like domain inhibits phosphotransfer from CheA-P to CheV and stabilizes CheV-P from autodephosphorylation.^49^ Our findings suggest that these effects are more likely the result of steric effects than conformational coupling between the two domains of CheV. The determination of CheV structures in the presence and absence of the stable phosphoryl group analog BeF_3_^-^ would be highly informative.
- Alexander *et al.*^15^ and Ortega & Zhulin^46^ proposed that the phosphorylation site in the receiver domain of CheV might act as a phosphate sink, while the CheW-like domain localizes CheV appropriately. Our results are consistent with such a hypothesis because no interaction between the two domains of CheV is required that might lead to divergence of CheW-like domains in CheW and CheV proteins. In a phosphate sink mechanism, when environmental circumstances change and phosphoryl groups stop flowing into the system from ATP, phosphoryl groups flow backwards from the CheY response regulator to CheA to the sink and are lost to hydrolysis. Localizing the sink near CheA would enhance efficiency. We note that another possible role of CheV is as a reservoir that retains phosphoryl groups in a buffering capacity.
- In a more dynamic role, phosphorylation-mediated dimerization of the receiver domain of CheV might control the ability of the CheW-like domain to bind with receptor and/or kinase partners. To the best of our knowledge, the multimeric state of CheV and any effect of phosphorylation on multimeric state have not been investigated.

Second, five (CheA.III, CheA.IV, CheA.V, CheA.VII, CheA.X) of the 12 CheA *Architectures* in Figure 2 contained a receiver domain immediately C-terminal to a CheW-like domain, which is reminiscent of CheV proteins. However, the fact that CheW-like domains in CheV and CheA proteins were in different *Classes* (1 vs. 3, 4, or 5) strongly suggests that adjacent CheW-like and receiver domains in CheA proteins do not act as CheV analogs. Consistent with this prediction, the adjacent CheW-like domain did not affect autodephosphorylation of the receiver domain from *Azorhizobium caulinodans* CheA,^51^ in contrast to *B. subtilis* CheV.^49^

Third, as far as we know, obvious questions about CheW-lineage proteins with multiple domains have not been explored experimentally. It is not known if all CheW-like domains of multi-domain CheW proteins are required for function, or if there are differences in functional roles between the domains. The fine structure of the data in Table 1 (specifically the number of sequences in a *Cluster* varied substantially as a function of *Context* within CheW.II and CheW.III *Architectures*) strongly suggests that the domains in a single multidomain CheW protein often belong to different sequence *Clusters* or even *Groups*, which could suggest domain specialization. On the other hand, because experimentally characterized single domain CheW proteins belonged to *Class* 1, which also includes ∼93% of domains in multi-domain CheW proteins, our results suggest that there is likely little to be learned about general CheW function by exploring multi-domain CheW proteins. Presumably, multi-domain CheW proteins support formation of complexes with specific components. This conjecture is supported by the observation that CheW.III proteins form cytoplasmic double layered arrays with CheA and a special type of double domain chemoreceptor in *Vibrio cholerae*.^52^

Approximately 1% of CheW-like domains in CheW proteins were CheA-like (CheW.IC *Type*) and formed a *Class* of their own. Some members of *Class* 2 appeared to span the gap between CheA proteins and CheW/CheV proteins (Figures 4 and S2-S25). The CheW-like domain is the most recently evolved fold in proteins that support bacterial chemotaxis.^44^ The birth of CheW-like domains allowed the separation of receptor and kinase functions into different proteins, in turn facilitating the integration of multiple stimuli sensed by arrays of different receptors into a single kinase. We are unaware of any evidence establishing whether CheW-like domains evolved first in CheW, CheA, or CheV proteins. Because CheV proteins are less common and less widely distributed than CheA or CheW proteins,^44^ it seems reasonable that CheV proteins came after CheA and CheW. Because members of *Class* 2 segregated with *Class* 1, *Class* 3 and in between in our analyses (Figure 4), *Class* 2 may represent a “missing link” in the evolution of CheW-like domains from CheW to CheA or from CheA to CheW. Closer examination of sequences in *Class* 2 may provide new insights into the evolution of chemotaxis systems.

CheW-like domains in most CheA proteins with one CheW-like domain were similar (*Class* 3), even though there were 10 different major *Architectures* for such proteins, including one (CheA.IX) that lacked a phosphorylation site (Hpt domain) and two (CheA.XI, CheA.XII) that lacked ATP binding sites (HATPase_c domain). Such “split” CheA *Architectures* are not degenerate, but can work together to reconstitute a functional unit.^53^ N-terminal CheW-like domains in CheA *Architectures* with two CheW-like domains (CheA.V.1, CheA.VI.1) were similar to CheW-like domains in CheA *Architectures* with a single CheW-like domain. The existence of *Class* 3 suggests that what is known about CheW-like domains in experimentally well-characterized CheA proteins is likely to be applicable to the majority of CheW-like domains in CheA proteins. The exceptions would be *Class* 4 and 5 CheW-like domains.

*Class* 4 CheW-like domains were found in CheA.VII and CheA.X *Architectures*, which have three Hpt domains and a C-terminal receiver domain. There were many minor CheA-lineage *Architectures* that contained up to 10 N-terminal Hpt domains and a C-terminal receiver domain. To the best of our knowledge, the only such protein to have been experimentally characterized is *Pseudomonas aeruginosa* ChpA, which controls Type IV pili rather than flagella. ChpA has six Hpt domains and supports an intramolecular phosphorelay.^54^ Phosphotransfer within ChpA proceeds from ATP to the Hpt domains close to the HATPase_c domain to the receiver domain to Hpt domains further from the HATPase_c domain. Thus, a plausible hypothesis is that a distinct *Class* of CheW-like domains is required to support intramolecular phosphorelays in CheA proteins with multiple Hpt domains and a receiver domain. Perhaps the receiver domain (immediately adjacent to the CheW-like domain) must be held in a particular orientation to be accessible to multiple Hpt domains.

*Class* 5 featured the C-terminal CheW-like domain in CheA proteins with dual CheW-like domains. *A. caulinodans* CheA,^51^ *Azospirillum brasilense* CheA1,^55^ and *Rhodospirillum centenum* CheA1^56^ have been explored experimentally and all belong to the CheA.V *Architecture* with two CheW-like domains. In each case, the C-terminal receiver domain is functionally important. However, the only experimental investigation of a *Class* 5 CheW-like domain of which we are aware was in *A. caulinodans* CheA, where removal of the C-terminal CheW-like domain diminishes chemotaxis but does not reveal the functional role of the *Class* 5 domain.^51^ *Class* 5 CheW-like domains were substantially different from experimentally characterized CheW-like domains. Therefore, because investigation of *Class* 5 is especially likely to reveal new information, *Class* 5 arguably represents the highest priority for future research on CheW-like domains.

*Class* 6 was a *Type* (CheW.IA) encompassing ∼20% of CheW proteins that appeared different from most CheW proteins (*Class* 1) in some analyses and similar in others. *Class* 6 CheW-like domains appeared to be different from the rest by *k*-medoid analysis (see Figure 3/Table 1). *Class* 6 appeared indistinguishable from *Class* 1 in the metric MDS analyses of random stratified subsampling in Figure 4 and Figures S2-S13, and the differences were subtle in the SimpLogo of Figure 5. *Class* 6 showed some separation from *Class* 1 in the non-metric MDS analyses of random stratified subsampling in Figures S14-S25. Method-dependent classification suggests that *Class* 6 is unlikely to represent a high priority for investigation. However, a companion manuscript reports additional analyses that may shed light on *Class* 2 and *Class* 6 CheW-like domains.^57^

### 4.2 Generally applicable methods to analyze protein domains as a function of architectural context

We have described a workflow (Figure 1) to analyze variation between sequences of a single protein domain as a function of architectural context within proteins. To ensure a robust workflow, we tested various approaches to optimize our method, including using multiple different amino acid substitution models and multidimensional scaling techniques to estimate and visualize relevant domain sequence subpopulations. Furthermore, the similarity of our partition results using these different models was compared using ARI and NMI analyses. We first applied a basic *k*-medoid clustering step to quantify the relative homogeneity of a given domain sequence set. This allowed us to identify constituent strata for representation in a more complete sequence analysis utilizing a random stratified sub-sampling scheme. We demonstrated our approach using CheW-like domains as a general case study. The results were largely independent of methods chosen, suggesting robust conclusions. Grouping sequence *Types* of CheW-like domains into six major *Classes* led to a variety of useful observations, insights, hypotheses, and suggested areas of future experimental investigation.

It might appear challenging to apply our approach to domains found in many distinct domain architectures. However, domain architecture distributions tend to have a long tail, with only a few organizations representing the majority of cases^58^. Here, 16 out of 111 architectures in Pfam that contained CheW-like domains incorporated ∼94% of CheW-like domains. Of course, individual “rare” architectures of interest could always be manually included with minimal difficulty. Another concern could involve the number of sequences to analyze. Our case study involved analysis of ∼8500 sequences. Significantly more or less sequences may be available for other domains. However, our approach should still be practical. The number of starting sequences to be analyzed can be managed by choice of an appropriate input data set. We chose (and recommend) the use of Representative Proteome sets as a starting point, where the sequence space size and granularity can be modified by changing the co-membership threshold of the relevant RP set cutoff (we used 35% as our threshold, but RP sets generated using 15%, 55%, and 75% are also available).^17^ The subsampling used in our analysis (see Figure 4) is a standard strategy to deal with impractically large data sets.

Finally, we developed and have made available (see Methods) SimpLogo, a new means of visualizing amino acid sequence conservation patterns across many subtypes of the same domain and for sequences of arbitrary length. Comparison of the SimpLogos in Figure 5 and traditional sequence logos in Figure S26 quickly reveal the utility of SimpLogos. Differences in gap patterns are immediately obvious. Identifying differences in amino acid composition at individual positions is more time consuming, but feasible. Traditional sequence logos are useful to communicate information about reasonably conserved positions but of little utility for most other positions, whereas SimpLogo provides useable information about all aligned positions within a given sequence.

## ACKNOWLEDGEMENTS

We thank Emily N. Kennedy and Sarah A. Barr for their insightful input and helpful discussions, including Sarah’s idea that receiver domains of CheV proteins might serve as reservoirs rather than sinks for phosphoryl groups.

This work was funded by National Institutes of Health grant GM050860 to Robert B. Bourret. The content is solely the responsibility of the authors and does not necessarily represent the official views of the National Institute of General Medical Sciences or the National Institutes of Health.

## Data Availability Statement

The data that support the findings of this study were derived from the following resource available in the public domain: Pfam (release v.33), http://pfam.xfam.org.

## Conflict of Interest Statement

The authors declare no conflict of interest.

## Notes

### Competing Interest Statement

The authors have declared no competing interest.

### Summary of Updates

Significant reorganization and clarification of presentation in response to peer review comments. No change in results.

