## Supplementary material for "Generalizable strategy to analyze domains in the context of parent protein architecture: A CheW case study": Text S1, Figures S1-S26

Revised for *Proteins: Structure, Function, and Bioinformatics*

5/31/22

### SUPPORTING INFORMATION

#### Text S1.

##### Gap patterns characteristic of CheW-like domain *Types*.

Figure 5 reveals obvious difference in gap patterns between different *Types* of CheW-like domains, which are summarized and interpreted below. All positions are described relative to *E. coli* CheW unless otherwise specified.

At positions 45, 55, and 56, the CheA-lineage *Types* generally exhibited sequence gaps, consistent with the observation that sequences of CheW-like domains from CheA proteins typically appeared shorter overall than from single domain CheW proteins when using the standard Pfam model (corroborated in manual CheW-like domain sequence alignments, data not shown). Experimental binding data suggest that positions 45-56 are involved in the interaction between the standalone CheW and CheA, at least in *E. coli.*[1,2] The CheW-like domain topology resembles that of a SRC Homology 3 (SH3) domain and is made up of two separate, pseudo-symmetric subdomains (1 and 2).[3] Subdomain 1 of the CheW-like P5 domain in *E. coli* CheA (approximately residues 514-531 and 596-640) interacts with subdomain 2 of standalone *E. coli* CheW (approximately residues 36-113; termed interface 1), primarily through the β3-β4-β5 regions of each individual subdomain (approximately residues 608-633 on P5 and 44-67 on CheW, based on homology modeling).[4] However, the complex signaling array/lattice formed by chemoreceptor:CheW:CheA oligomers is facilitated by the formation of a secondary interface between subdomain 1 of standalone CheW and subdomain 2 of P5 (approximately residues 114-134 on CheW and 539-556 on P5, based on homology modeling; termed interface 2), an interaction that contributes substantially to the cooperativity of the signaling array.[5] Positions 45-56 did not overlap with known chemoreceptor binding surfaces on CheA or CheW, implying that differences in gap character seen in Figure 5 likely affect the primary and/or secondary associations between the CheW-like domains of CheA- and CheW-lineage *Architectures* and not direct interactions with the chemoreceptor. Because subdomain 2 serves distinct purposes in CheA and CheW, differences in the corresponding functional residues seemed reasonable.

There were two exceptions to the described dichotomy between CheA-lineage and CheW-/CheV-lineage *Types* regarding gaps at positions 45, 55, and 56. First, *Types* CheA.V.1 and CheA.VI.1 exhibited only a partial gap pattern at position 45. Recall that *Types* CheA.V.I and CheA.VI.1 fell into *Cluster* H (Figure 3/Table 1). Although ultimately considered part of *Group* 1, *Cluster* H showed significant deviation from the rest of the CheA-lineage *Types*, with increased resemblance to *Group* 2 (*Cluster* P, made up of *Types* CheA.V.2 and CheA.VI.2 from the same *Architectures*) in Figure 3. However, *Cluster P* exhibited gaps at position 45 (Figure 4). Second, the gap patterns at positions 45, 55, and 56 exhibited by *Type* CheW.IC were more reminiscent of the CheA-lineage sequences than the rest of the CheW-lineage *Types*. This is consistent with observations that (i) the CheW.IC *Type* at least partially resembled CheW-like domains from CheA-lineages (both belonged to *Group* 1 in Table 1) and (ii) as *Class* 2 spanned the sequence "gap" between CheA (*Class* 3) and CheW (*Class* 1) regions of Figure 4.

CheA-lineage *Types* (*Classes* 3-5) exhibited stretches of gaps at positions 120-131, which corresponds to subdomain 1 in CheW-like domains. Therefore, sequence differences here could also affect the association between CheA- and CheW-/CheV-lineage proteins. CheV- and CheW-lineage *Classes* 1 and 6 shared the gap pattern near position 120 with CheA-lineage *Classes* 3-5 but lacked the stretches of gaps in subdomain 1 near positions 124 and 130. In contrast, CheW.IC (*Class* 6) again more closely resembled the gap profile seen in the CheA-lineage *Classes* 3-5 at positions 120-131.

*Type* CheA.V.2 and CheA.VI.2 domains (*Class* 5) displayed a slightly modified gap profile at positions 122-130 and exhibited a more diffuse level of gap character at positions just upstream of the region (indicated by subtly "shrunken" boxes stretching back to residue position 110). Although the role(s) played by positions 109-110 in array formation is unclear, structural and biochemical data suggest that residue 108 is directly involved in chemoreceptor binding.[6] In the dual CheW-like domain CheA.V and CheA.VI *Architectures*, the "first" position CheW-like domains (*Types* CheA.V.1 and CheA.VI.1) did not exhibit the same deviation in gap character at residues 110-119 as *Class* 5, suggesting that the "second" position CheW-like domains may bind chemoreceptors that are distinct from those of their N-terminal counterparts, or perhaps bind entirely different partners through this region.

**Residue identities characteristic of CheW-like domain *Types*.**

Careful inspection of Figure 5 revealed several characteristic deviations in residue frequencies at key positions between *Types*, many of which involved *Class* 5 domains. Again, all positions are described relative to *E. coli* CheW.

The entire region between positions 108-134 was filled with conserved charged and polar residues that are likely involved in intermolecular/interdomain interactions in CheA-lineage *Types* but that were absent in the other *Types*. At position 110, several of the CheA-lineage *Types* encoded a negatively charged (red) residue (particularly in CheA.II/III/IV/V.1/VI.1/VII/X), whereas the position was less conserved in the CheV- and CheW-lineage *Classes* 1 and 6. The gap character was increased at position 110 in *Class* 5. Based on our homology models and alignment against another crystal structure of dimeric *T. maritima* CheA (residues 290-671 relative to the *T. maritima* protein; PDB ID 1B3Q), residue 110 likely interacts with the helical dimerization domain of the CheA kinase, possibly explaining the change in conservation in CheA-related *Types* compared to the other lineages. Similarly, position 113 was strongly hydrophobic (green), position 114 was strongly positively charged (blue), and position 115 was polar/Pro (yellow/white) in almost all the CheA-lineage *Types* except *Class* 5, but not in the CheV- and CheW-lineage *Classes* 1 and 6. These positions are likely related to formation of the interface between the CheW-like domains of CheA and CheW.

*Class* 5 domains exhibited distinct patterns of charged amino acids. Position 38 was positively charged (blue) in *Class* 5 but negatively charged (red) in all other *Classes* except 4. *Class* 5 also exhibited a pair of conserved, negatively charged (red) residues at positions 40-41, a unique trait not exhibited by any other *Class*. In *E. coli,* the region equivalent to position 40 is involved in chemoreceptor binding,[1,2,6] again suggesting different sensor/partner specificity for *Class* 5 domains.

At position 49, all four of the CheW-like domains in the CheA.V and CheA.VI *Architectures* shared a conserved negatively charged (red) residue. The remaining CheA-lineage *Types* exhibited substantially lower conservation, leading to brownish coloration, whereas the CheV- and CheW-lineage *Types* primarily exhibited a conserved Pro residue (white). Equivalent residues in *E. coli* are involved in the association of CheW-like domains in CheA and CheW, based on the homology models.

*Class* 4 *Types* (CheA.VII and CheA.X) did not appear obviously different than *Class* 3 *Types* (most CheA *Types*) in Figure 5 except for Pro at position 87. The only *Type* that consistently stood out in *Classes* 3 and 4 was *Type* CheA.VIII. At position 27, most *Types* exhibited a low level of conservation, leading to a reddish-brown color. However, the CheA.VIII *Type* featured almost exclusively negatively charged (red) residue types. Closer examination of the traditional sequence logo and alignment for the CheA.VIII *Type* revealed a nearly invariant Glu. In our *E. coli* CheW homology model, the negatively charged Glu side chain of position 27 formed a hydrogen bond with the hydroxyl group of Tyr29 (brown) near the site of interface 2, which is essential for higher-order oligomer formation.

At position 41, there was an increased conservation of positively charged (blue) residues in many of the *Types* from CheA- and CheW-lineages, including CheA.V.1, CheA.XI, CheA.XII, CheA.VII, CheA.X, and to a lesser extent CheW.IB, CheW.III.1 and CheW.III.3. However, *Class* 5 domains exhibited an increase in negative charge (red) at the same position. *Type* CheA.VIII featured a highly conserved Gln (yellow) at position 41 followed by a similarly conserved Pro (white). The Gln Pro pair was not found in other *Types*. Experimental data and our homology models suggested that position 41 is involved in both CheA/CheW association and chemoreceptor binding.[2]

We did not observe any positions where *Class* 6 (CheW.IA), representing ~20% of CheW-lineage domains, differed from *Class* 1 (most CheW domains and CheV domains). However, we did observe some variation within *Class* 1:

- Position 56 exhibited gaps in most CheA-lineage *Types* and a strong hydrophobic (green) character in most CheW-lineage *Types* but exhibited increased negative charge (red) in CheA.VIII, CheV.I and CheW.III.3, and a positively charged ([blue) residue in CheW.IB. *In E. coli* CheW, experimental data indicate that position 56 is involved in the formation of interface 1, forming a hydrogen bond with Tyr81. Our homology models suggested that the region is likely substantially altered in CheA-lineage structures, as indicated by the consistent gaps at positions 55-56,[1] with the exception of CheW-like domains from *Type* CheA.VIII (which includes the P5 subdomain of *E. coli* CheA). The deviations seen at position 56 were likely a result (or cause) of differences in primary CheW/CheA association interactions between various combinations of CheA and CheW *Types*.
- At position 32, most *Types* exhibited Pro (white), except *Types* CheW.IB, CheW.II.1, CheW.II.2, and CheV.I, which featured increased negatively charged

(red) residues in the first three, and almost exclusively Asn (yellow) in the latter. Position 32 is proximal to the chemoreceptor binding interface (according to our homology model for the CheW-like domain of *E. coli* CheA) and is likely involved in the positioning and/or formation of corresponding intermolecular interactions. The high conservation of a Pro residue in most *Types* suggested an important structural feature.

- Position 74 featured a seemingly buried hydrophobic (green) residue in most CheW-like domains, with the exceptions of *Types* CheW.IB, CheW.II.1 and CheW.III.3, which instead displayed positively charged (blue) residues (primarily Lys). The entire region between positions 72-74 was significantly more positively charged in the CheW-lineage *Types*, with the possible exception of CheW.IA. The implications of this unique characteristic are unknown, because the positions are not known to be involved in receptor binding or interface formation.
- At positions 86-87 (a critical residue pair involved in chemoreceptor binding in *E. coli* CheW[1,6]), most of the CheW-lineage *Types* exhibited polar (yellow) and positively charged (blue) residues, except for CheW.II.1 and CheW.IC, which featured negatively charged/polar (red/yellow) and small-volume/hydrophobic (dark grey/green) residue pairs, respectively.

Finally, CheA.VI.1, CheW.II.2 and CheW.III.2 were the only *Types* with a strongly conserved Cys (cyan) residue, found at positions 24, 50-51, or 135 respectively. To the best of our knowledge, no naturally occurring disulfide bond formation has been reported involving a CheW-like domain, though conservation implied that the Cys residues were functionally or structurally significant in some way. The role of position 24 is currently unclear. Positions 50 and 135 are thought to be involved in the association of CheA- and CheW-lineage domains.

**Table S1. Functional CheW-like domain residues.** Putative functional residues for the *E. coli* CheA-P5 and CheW domains, obtained from literature sources and homology models described in this study.

**FIGURE S1. Multidimensional scaling plots showing representative mediods from each of the 21 *Clusters* generated using dissimilarity matrices calculated with the four designated substitution models.** Each clustering attempt reconstitutes roughly the same data structure, consisting of three main cluster *Groups* (numbered 1, 3 or 4 in the top left panel, GONNET) along with a weaker, more intermediate *Group* (numbered 2 in the top left panel, GONNET) intersecting *Groups* 1 and 3. Although the colors and specific cluster designations may shift between analyses (overall cluster sizes are slightly altered depending on the substitution matrix used, affecting *Cluster* names), the actual partitioning of CheW sequences was highly consistent, regardless of the substitution model used (Adjusted Rand Indices and Normalized Mutual Information scores were 0.6-0.7 and 0.7-0.8, respectively, indicating highly similar clustering behavior).

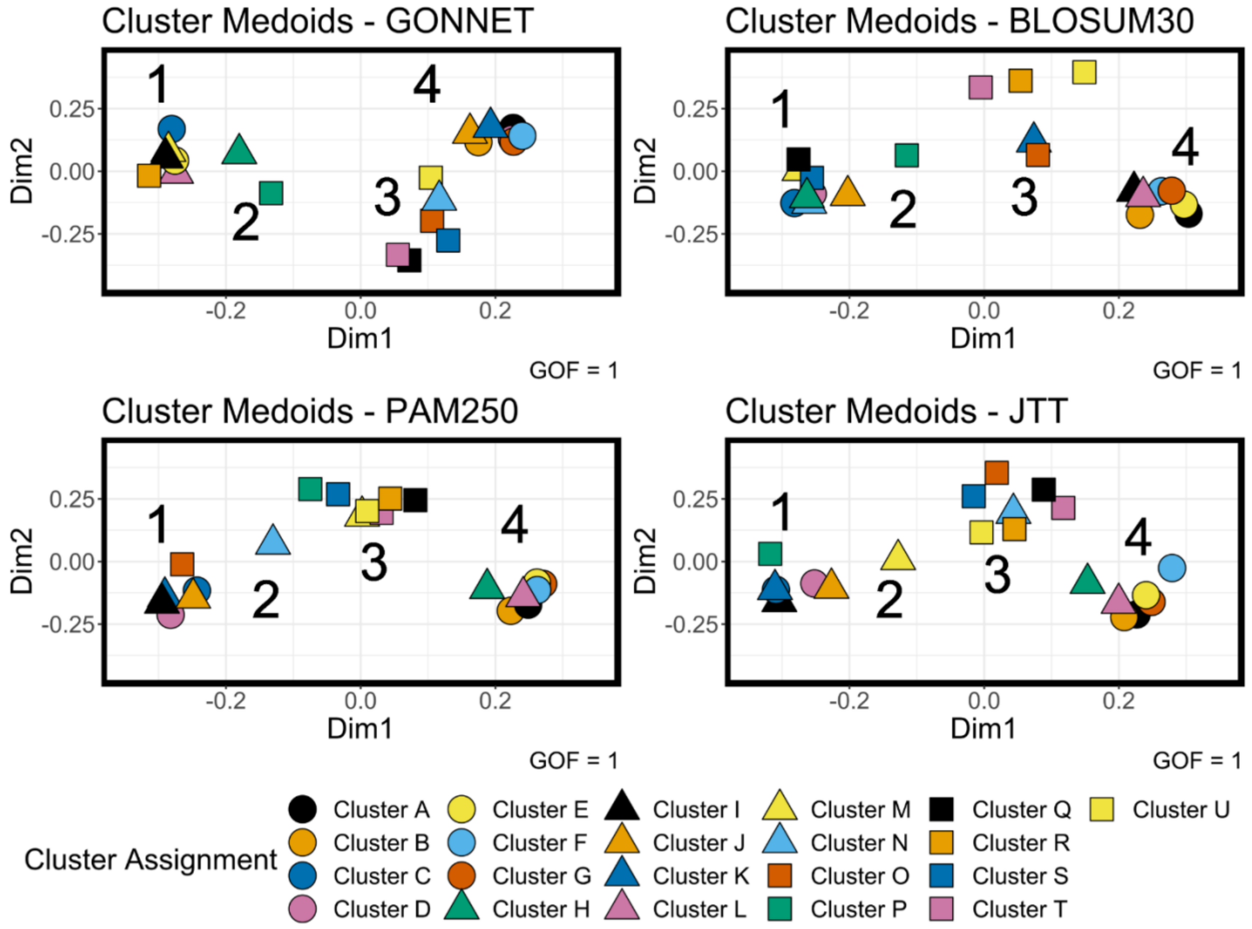

**FIGURES S2-S13.** Remaining metric MDS (mMDS) solutions produced for each of the subsampling iterations described in the Results section (using dissimilarities generated with the corresponding sustitution matrix; GONNET, JTT, BLOSUM30 and PAM250). Despite arbitrary rotations and reorientations of the coordinates, each solution demonstrates similar results. Dim1, Dim2, Dim3 = Dimensions 1, 2 3.

**FIGURE S2. GONNET-derived mMDS solutions – Iterations #1-4**

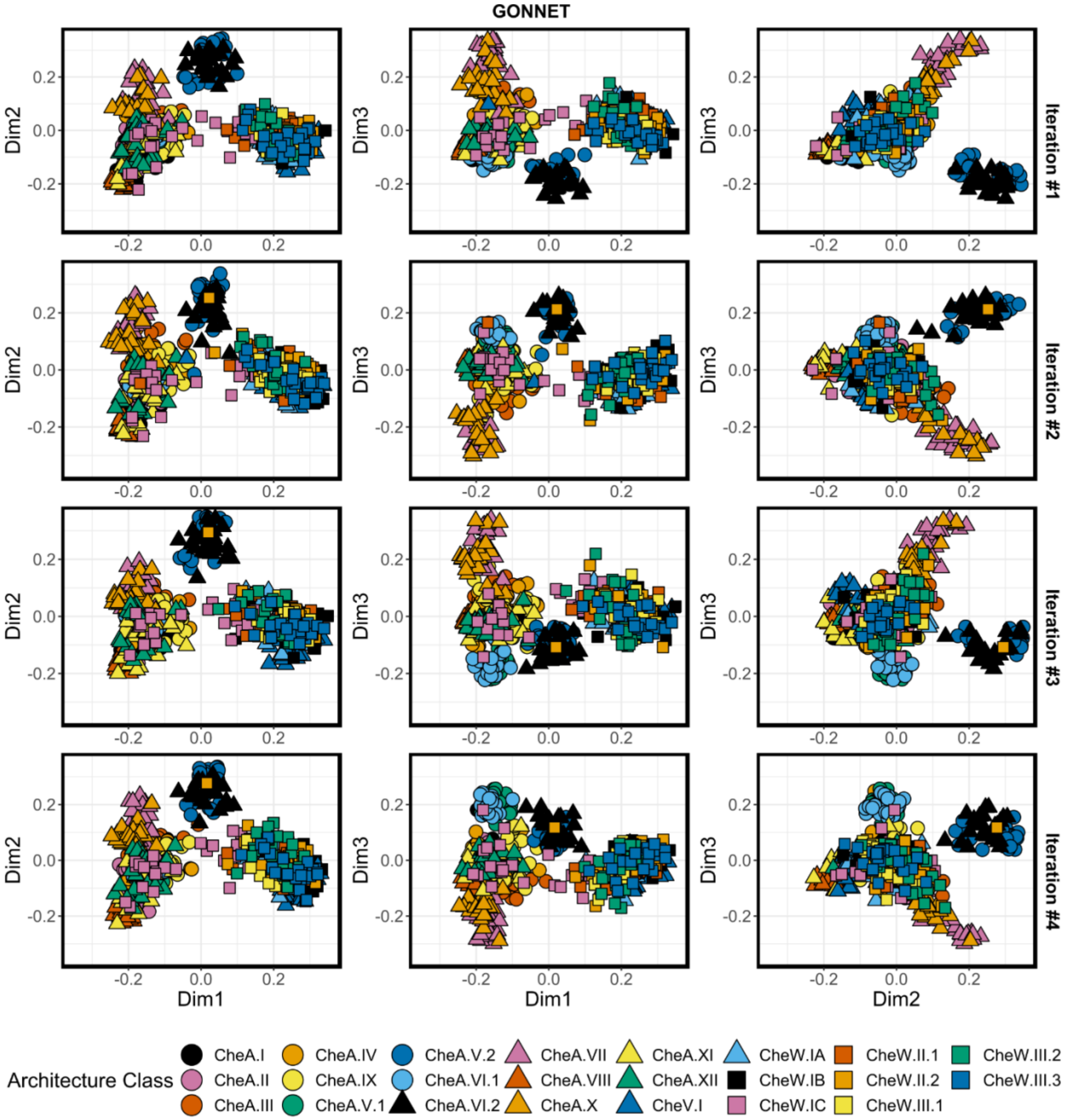

#### FIGURE S3. GONNET-derived mMDS solutions – Iterations #5-8

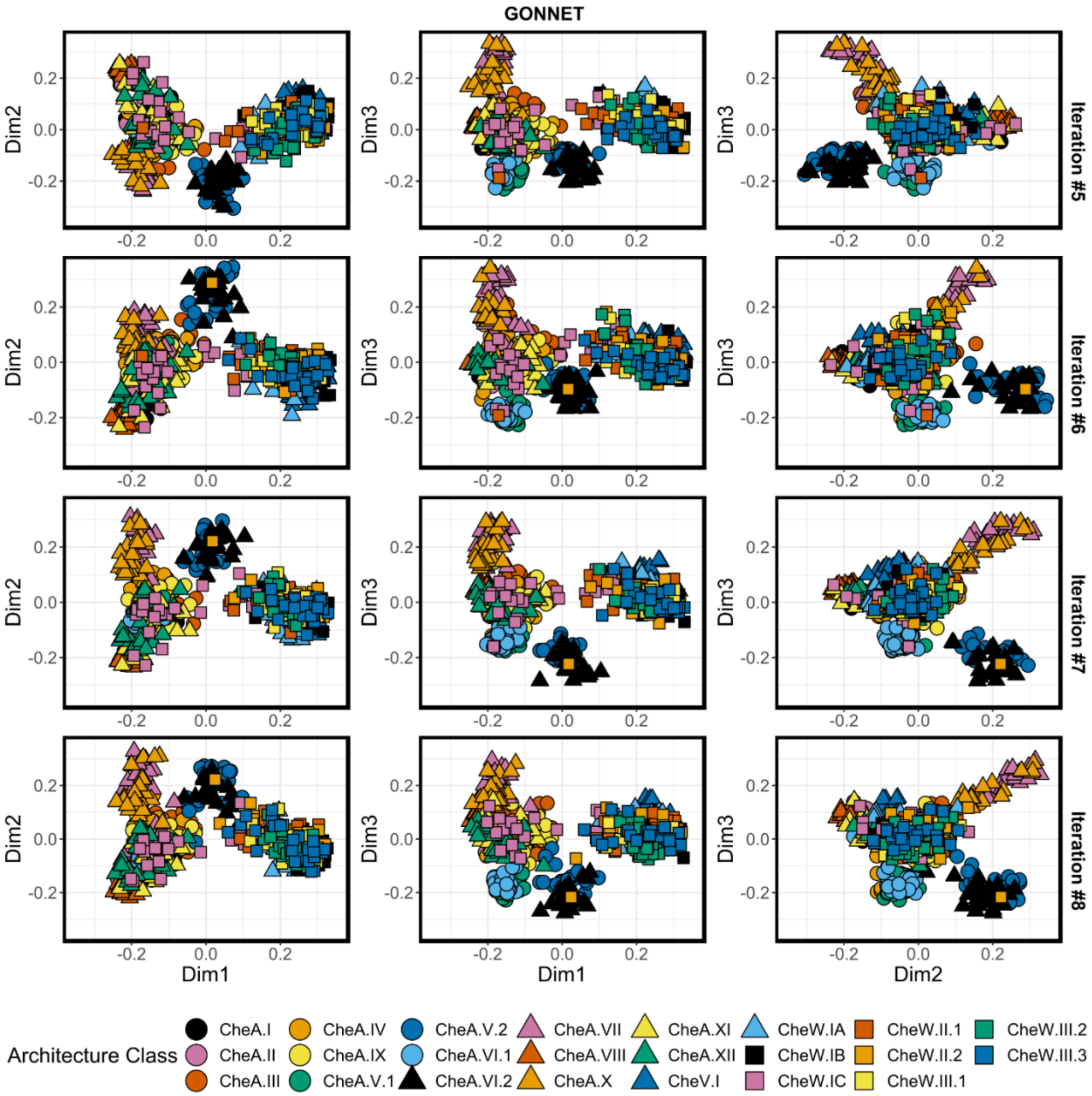

**FIGURE S4. GONNET-derived mMDS solutions – Iterations #9-10**

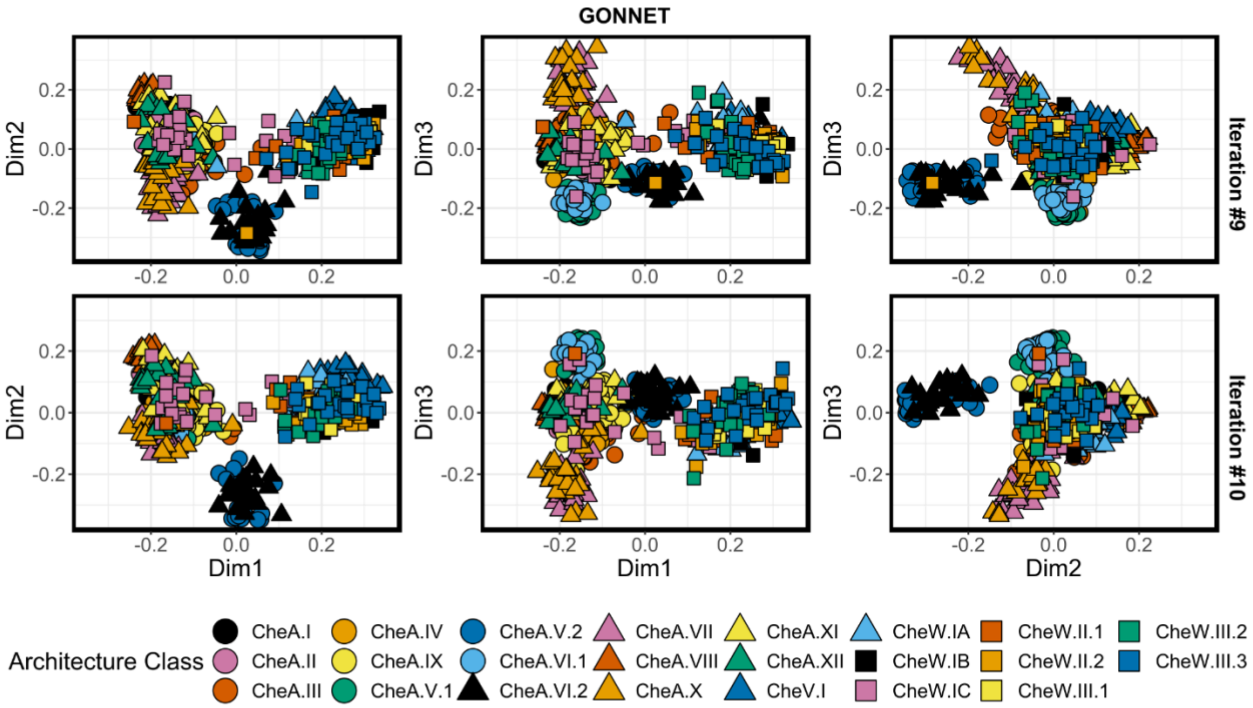

**FIGURE S5. JTT-derived mMDS solutions – Iterations #1-4**

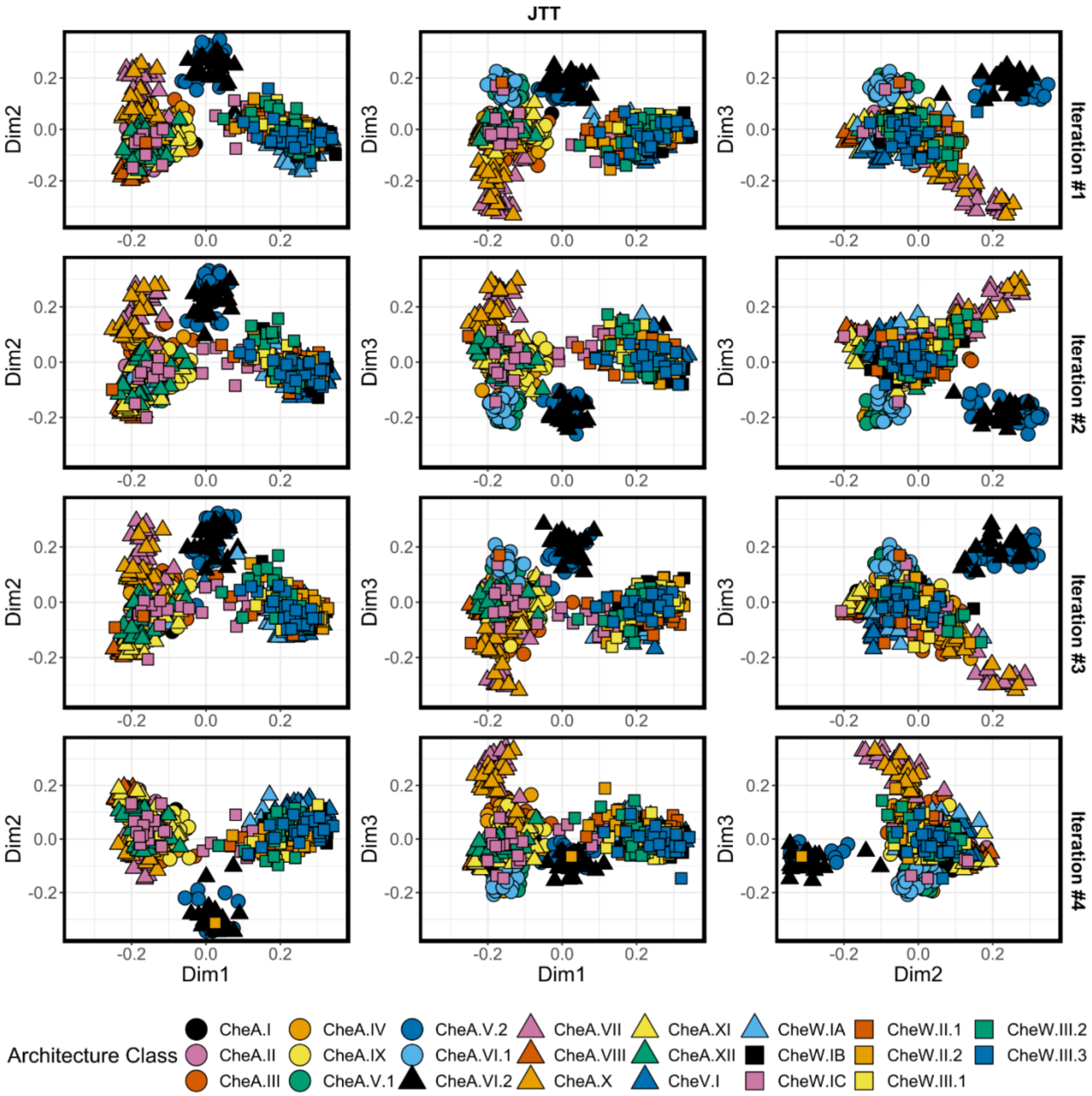

**FIGURE S6. JTT-derived mMDS solutions – Iterations #5-8**

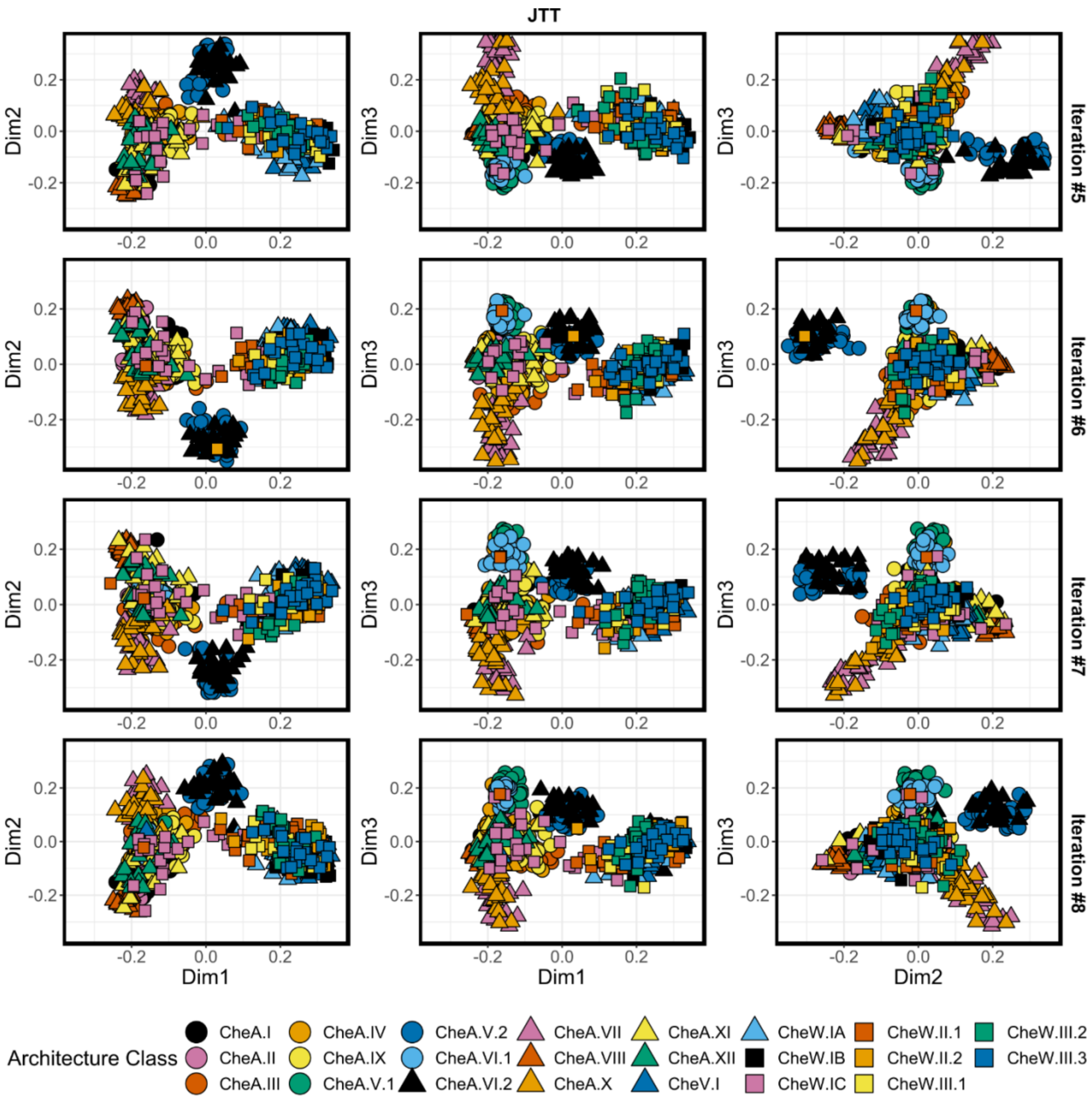

**FIGURE S7. JTT-derived mMDS solutions – Iterations #9-10**

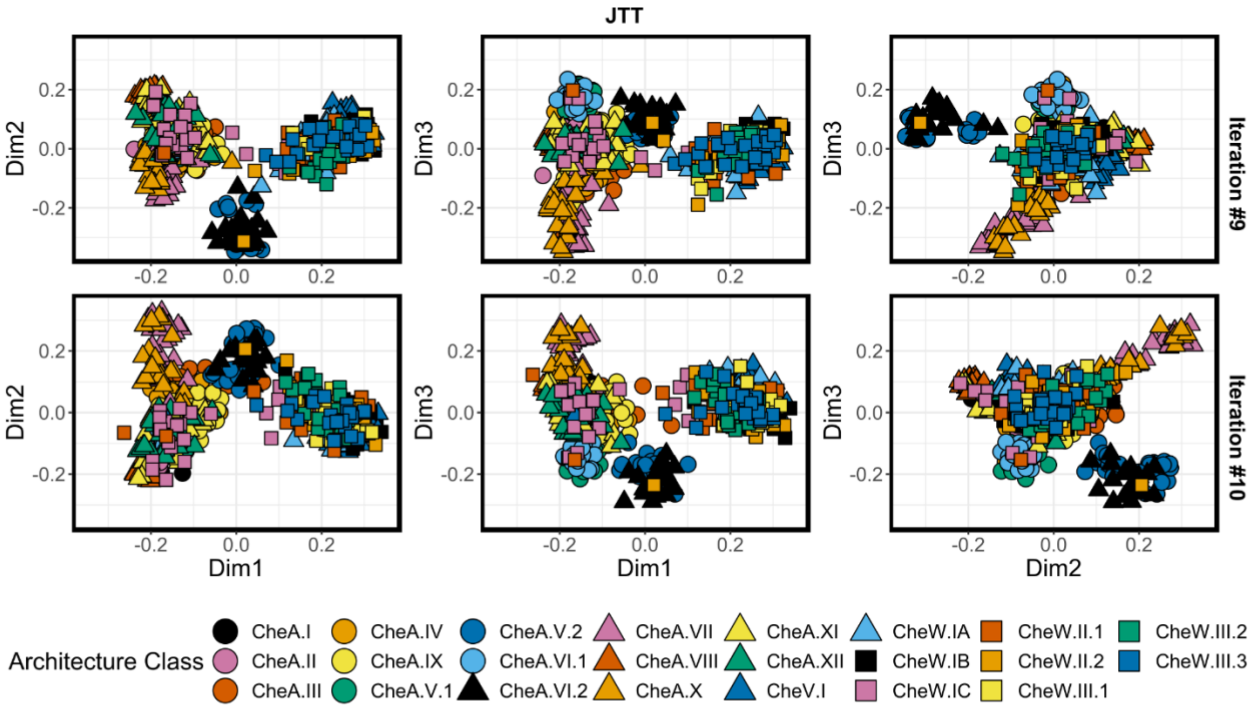

**FIGURE S8. BLOSUM30-derived mMDS solutions – Iterations #1-4**

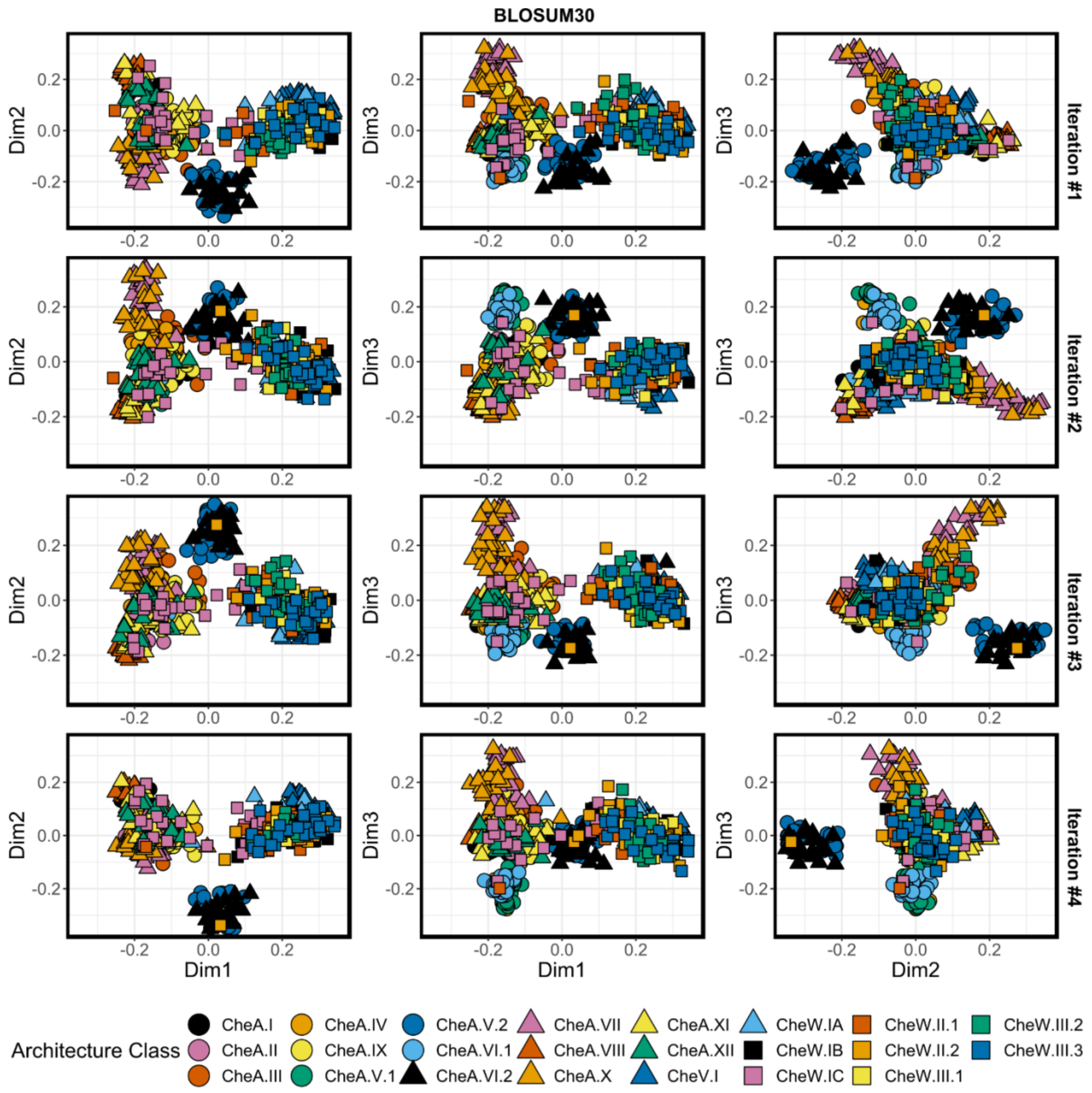

**FIGURE S9. BLOSUM30-derived mMDS solutions – Iterations #5-8**

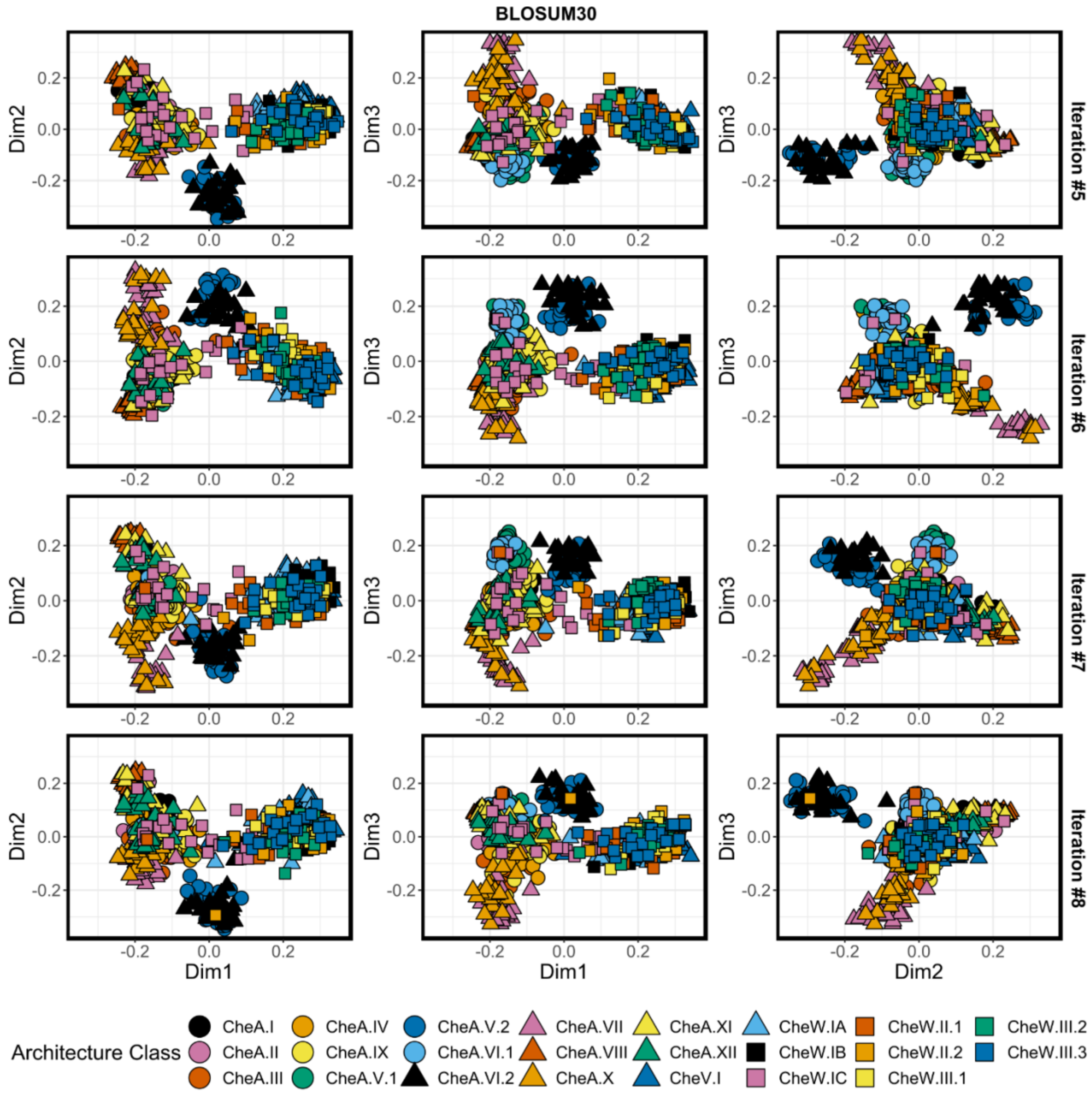

**FIGURE S10. BLOSUM30-derived mMDS solutions – Iterations #9-10**

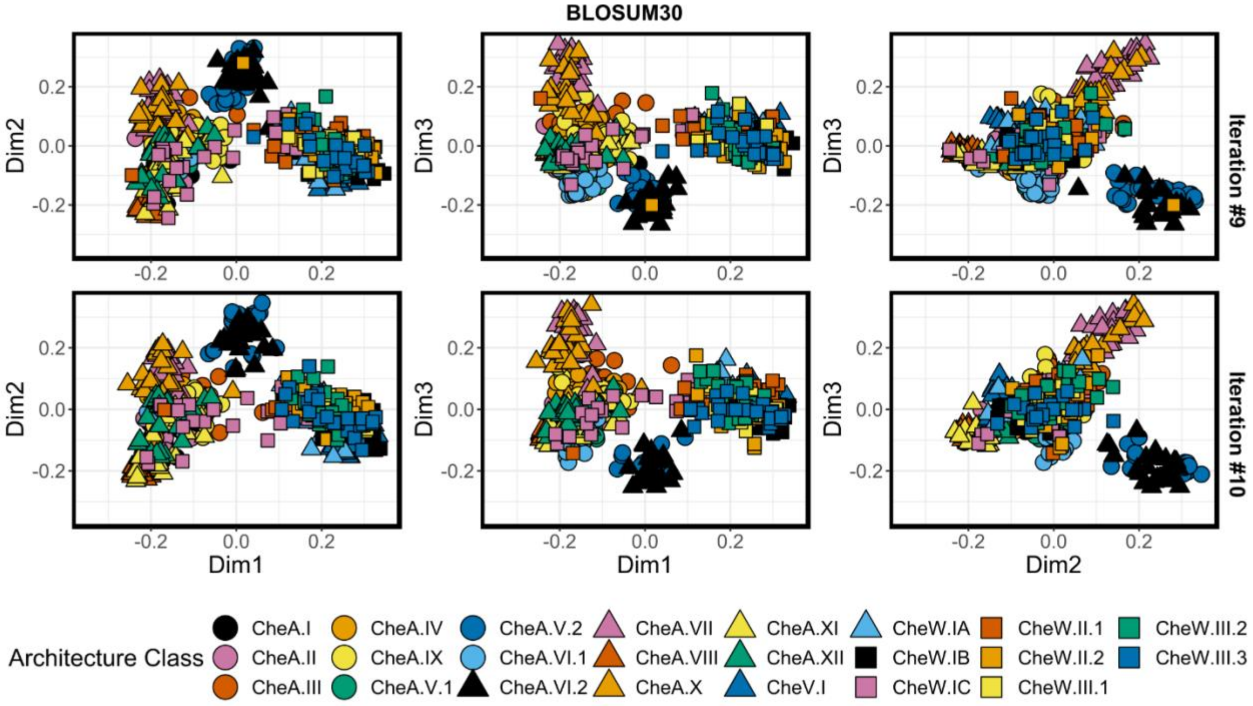

**FIGURE S11. PAM250-derived mMDS solutions – Iterations #1-4**

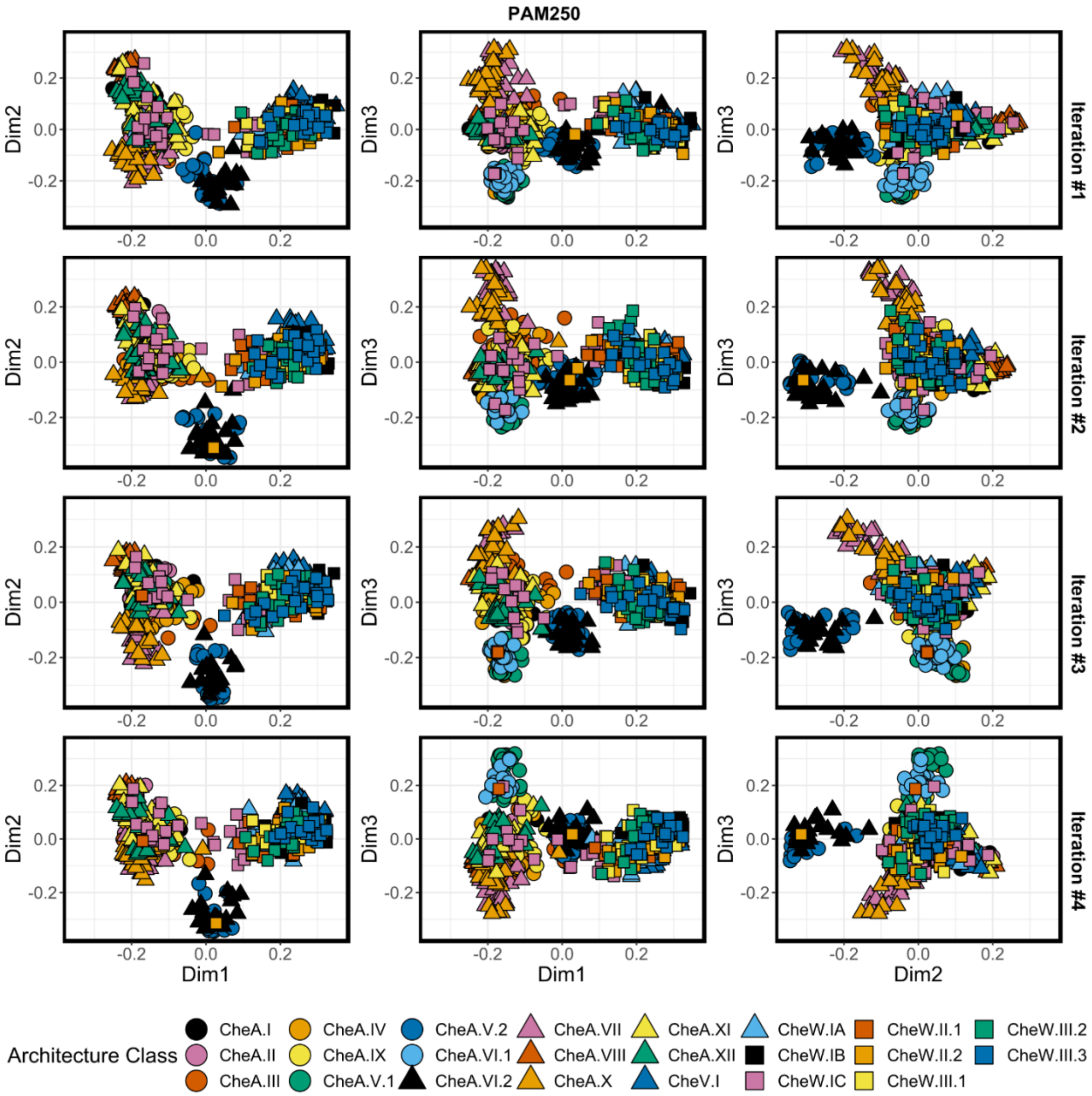

**FIGURE S12. PAM250-derived mMDS solutions – Iterations #5-8**

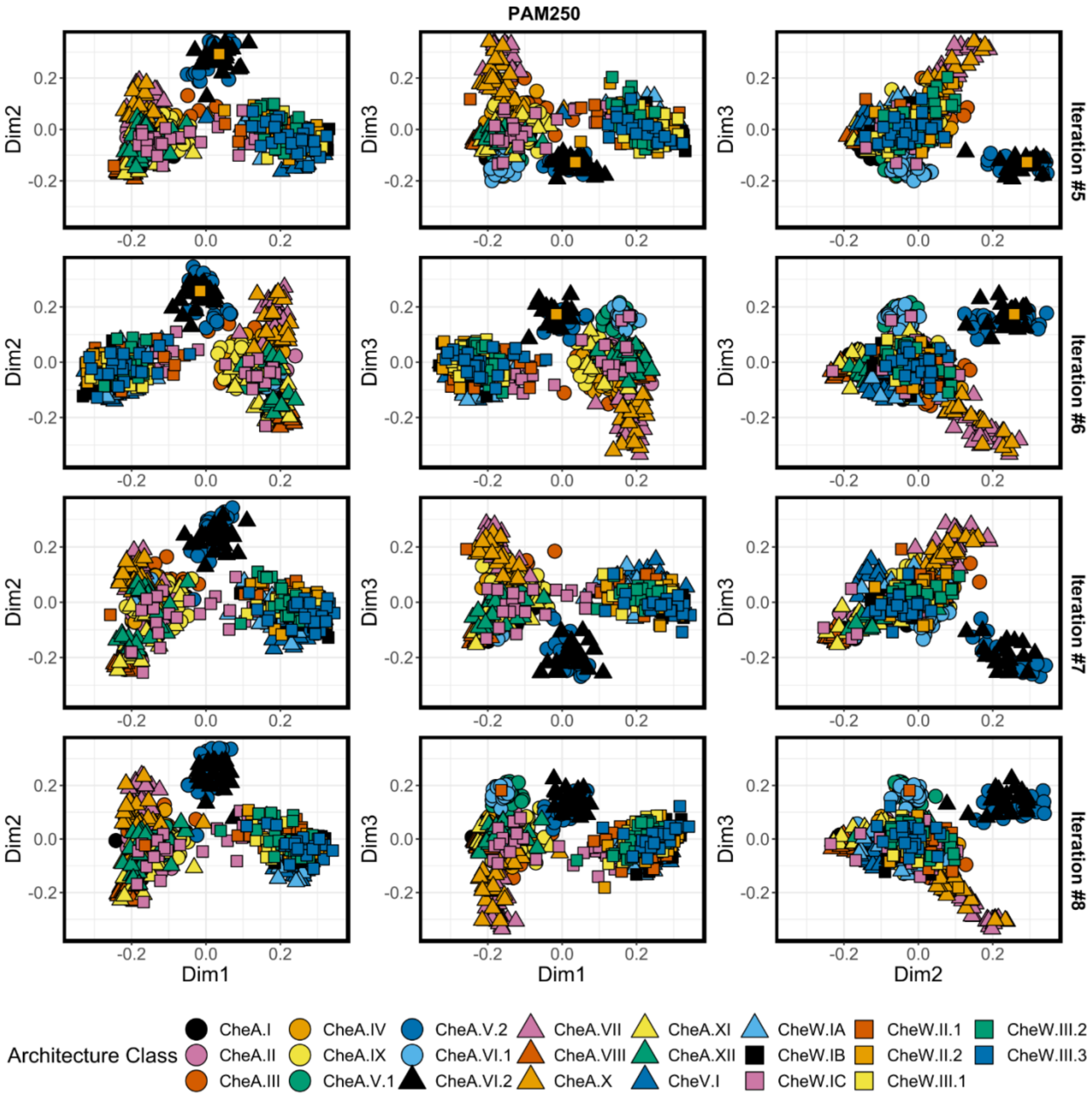

**FIGURE S13. PAM250-derived mMDS solutions – Iterations #9-10**

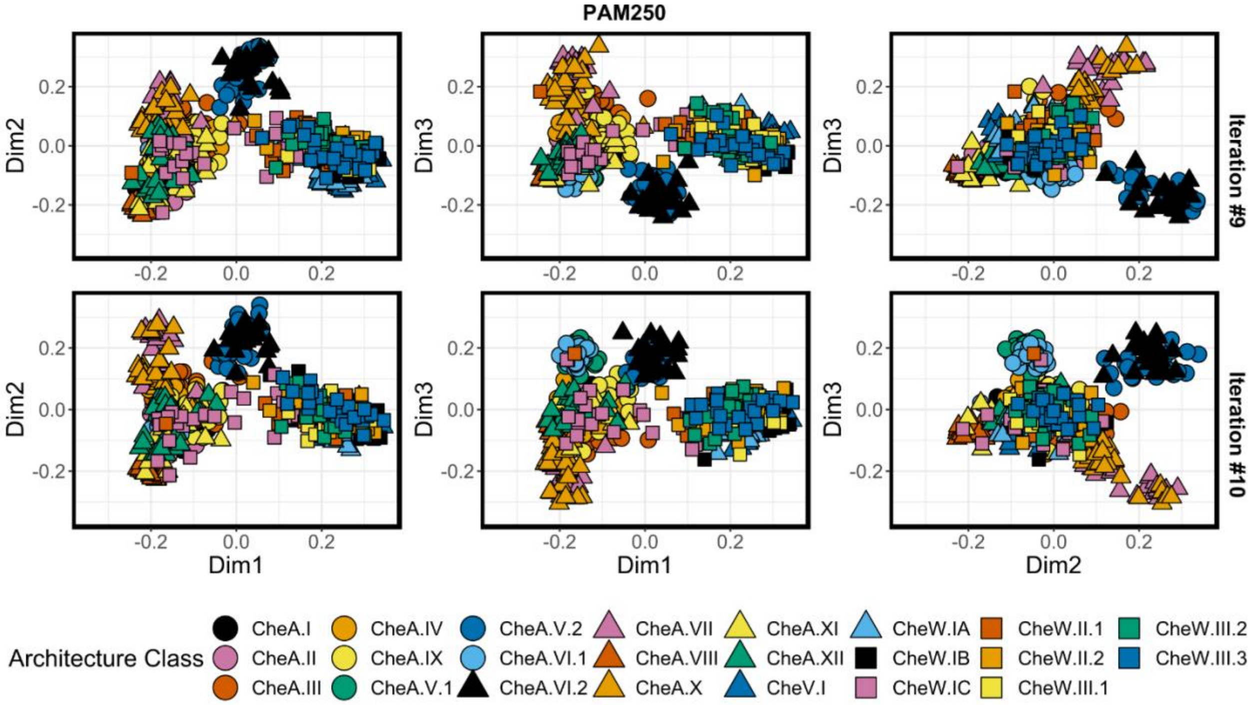

**FIGURES S14-S25.** Remaining non-metric MDS (nMDS) solutions produced for each of the subsampling iterations described in the Results section (using dissimilarities generated with the corresponding sustitution matrix; GONNET, JTT, BLOSUM30 and PAM250). Despite arbitrary rotations and reorientations of the coordinates, each solution demonstrates similar results. Each solution is also highly reminiscent of the metric MDS-derived layouts.. Dim1, Dim2, Dim3 = Dimensions 1, 2 3.

**FIGURE S14. GONNET-derived nMDS solutions – Iterations #1-4**

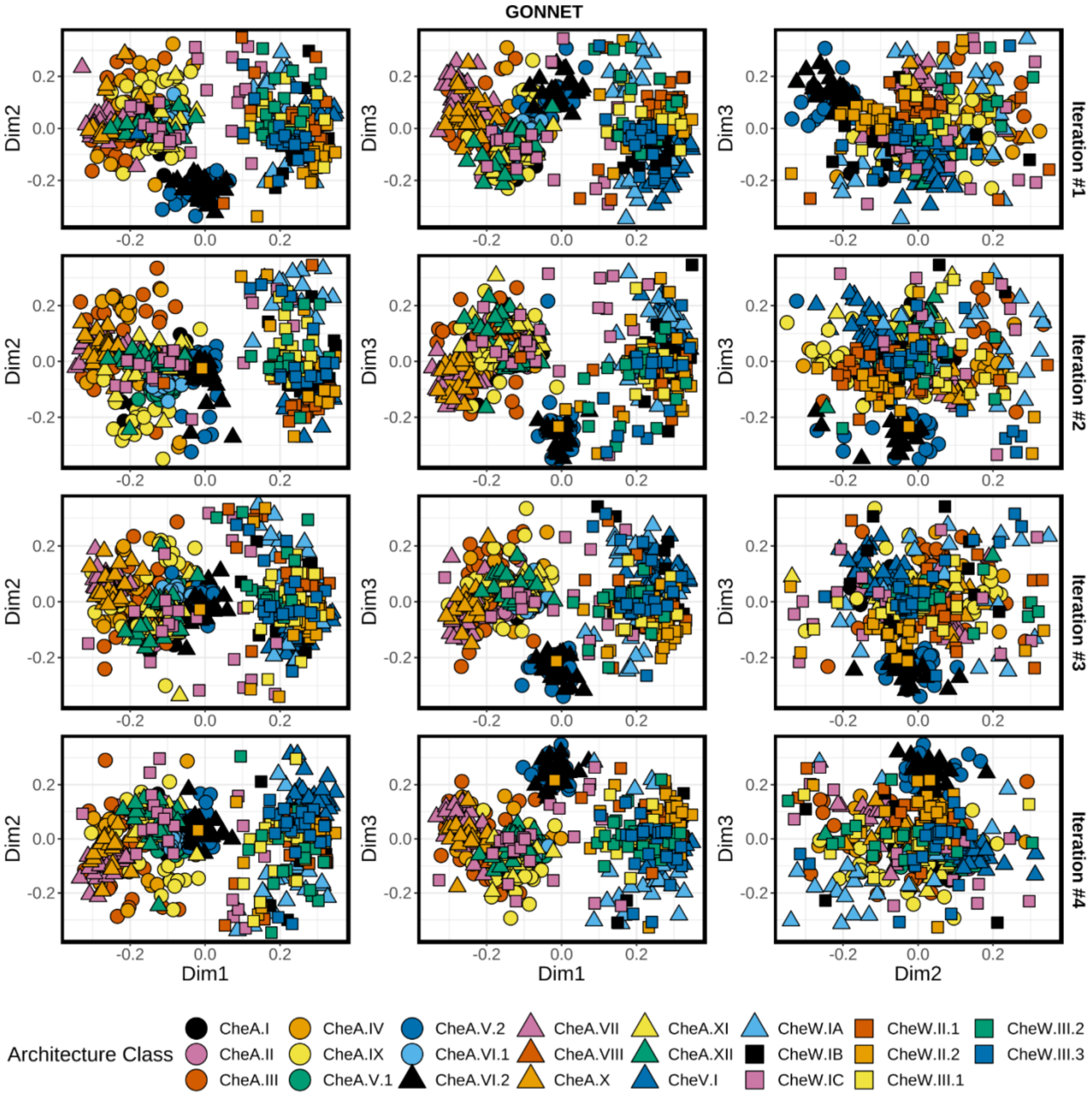

**FIGURE S15. GONNET-derived nMDS solutions – Iterations #5-8**

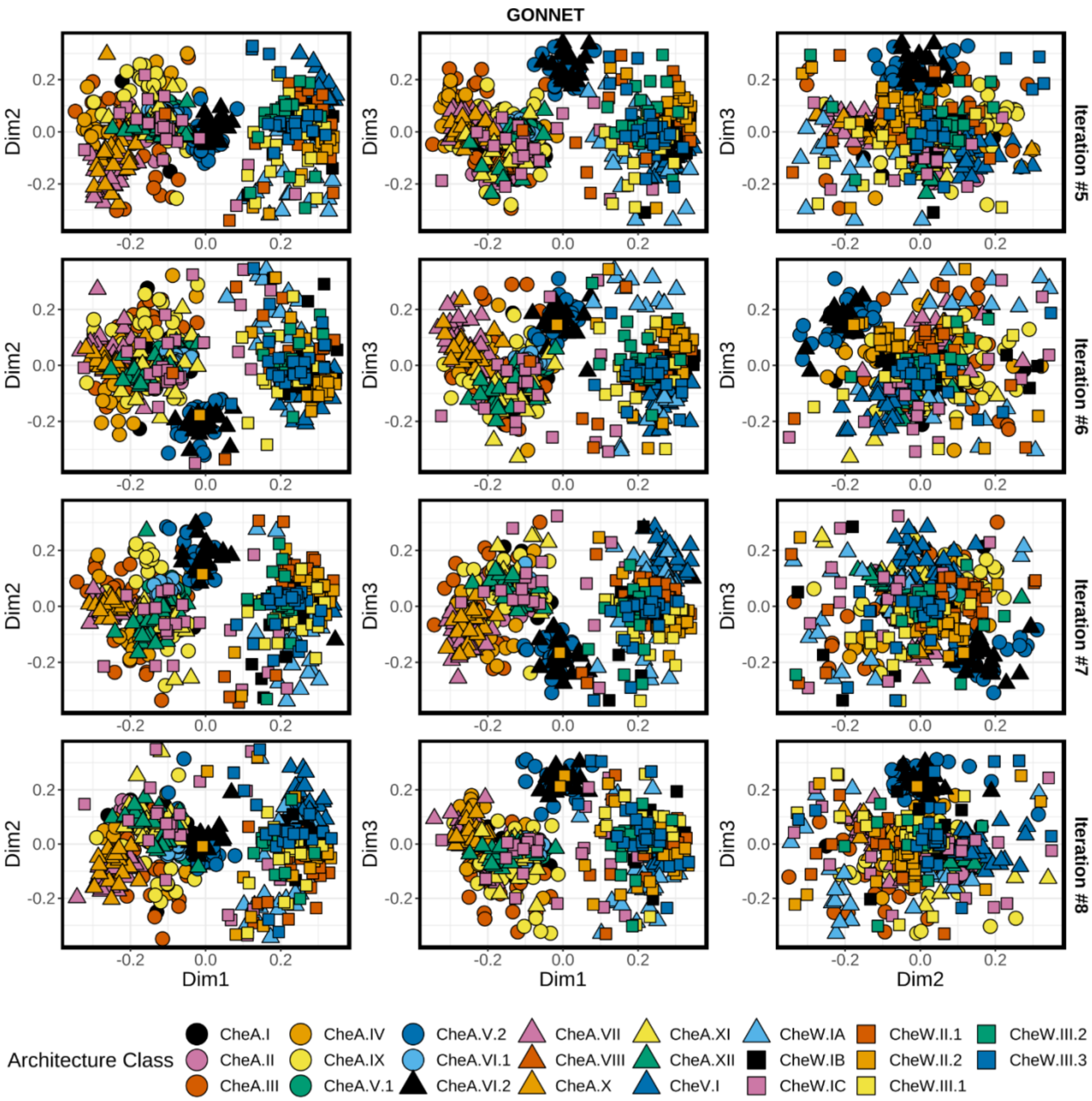

**FIGURE S16. GONNET-derived nMDS solutions – Iterations #9-10**

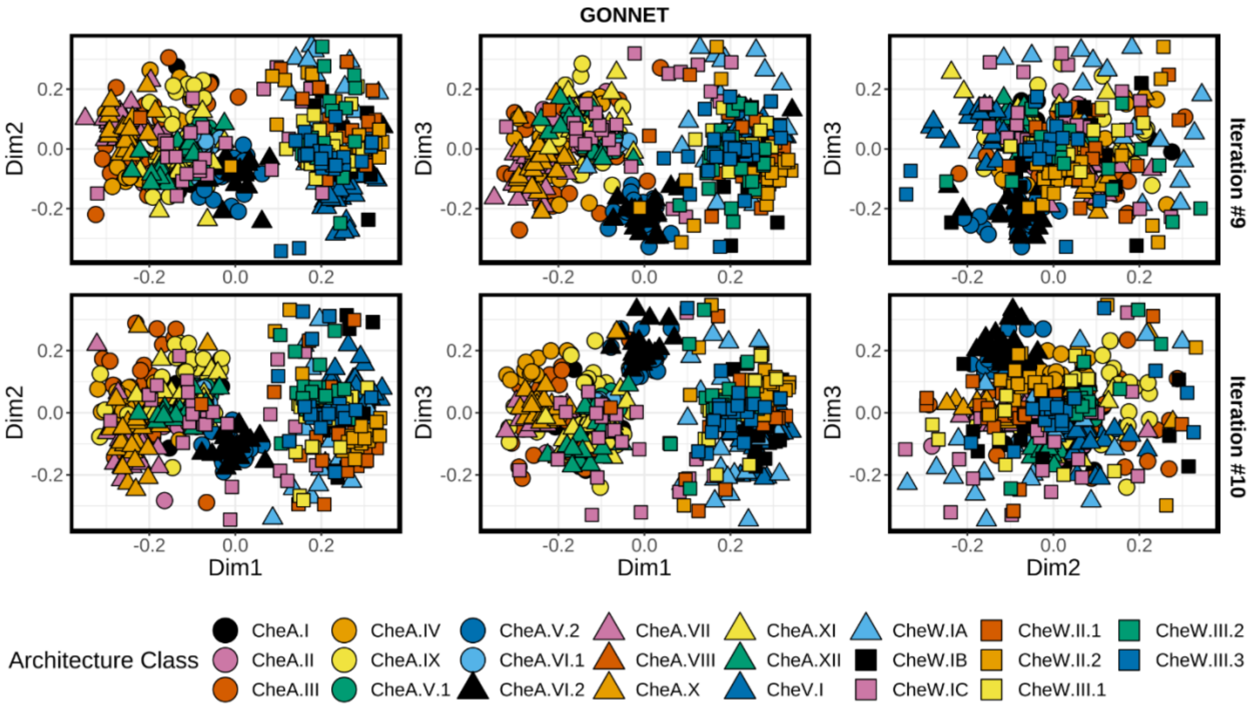

#### FIGURE S17. JTT-derived nMDS solutions – Iterations #1-4

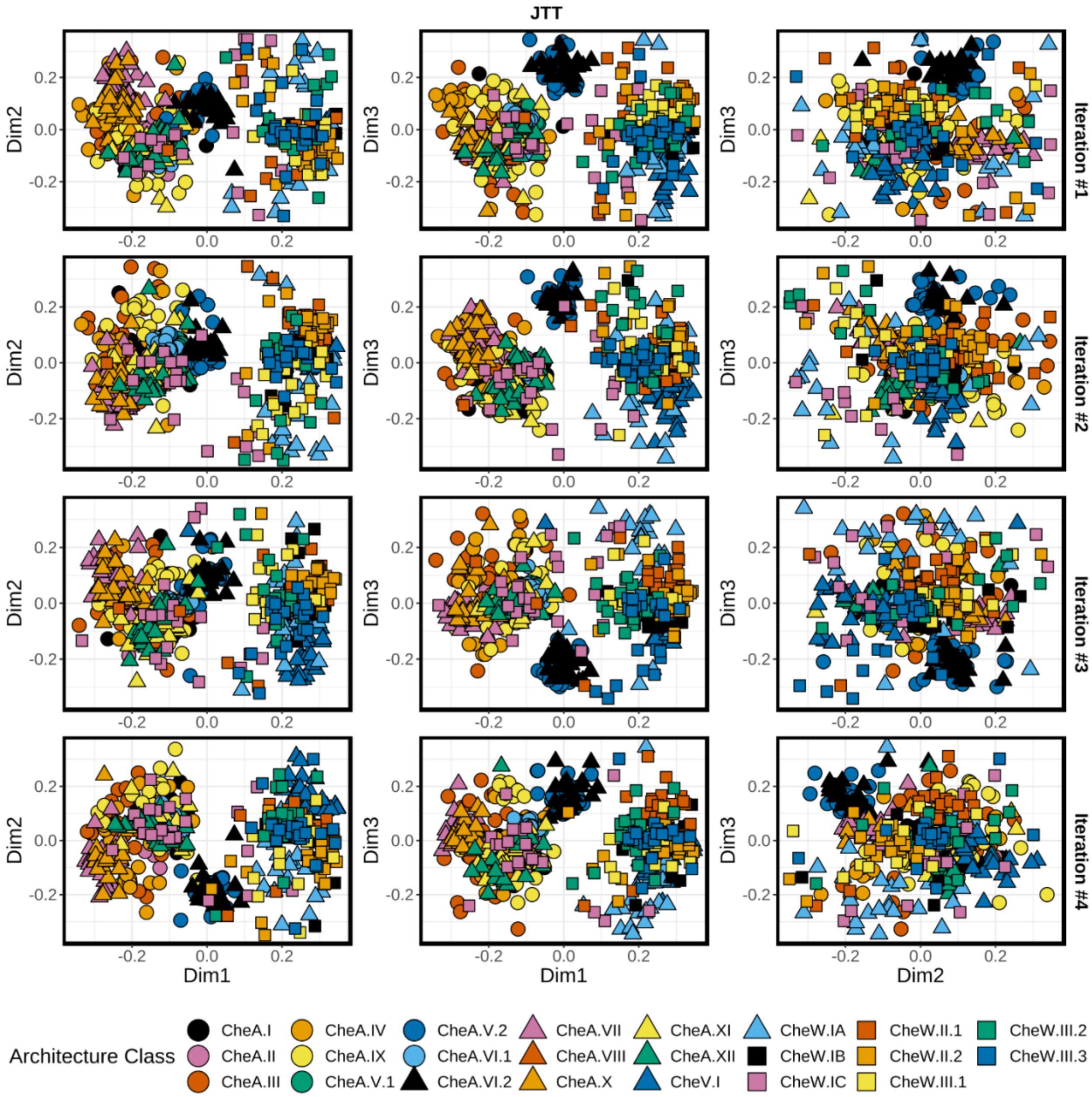

**FIGURE S18. JTT-derived nMDS solutions – Iterations #5-8**

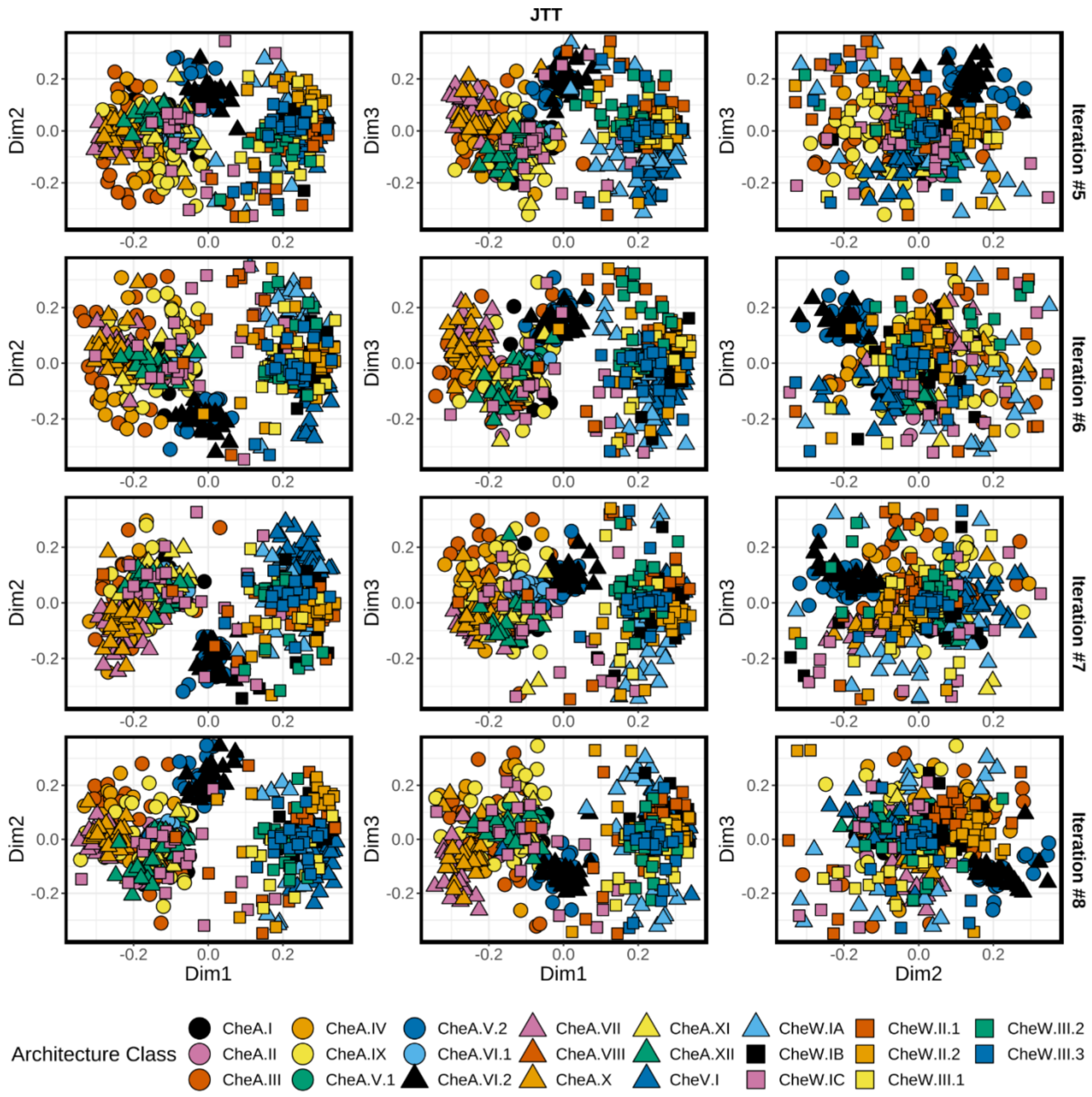

**FIGURE S19. JTT-derived nMDS solutions – Iterations #9-10**

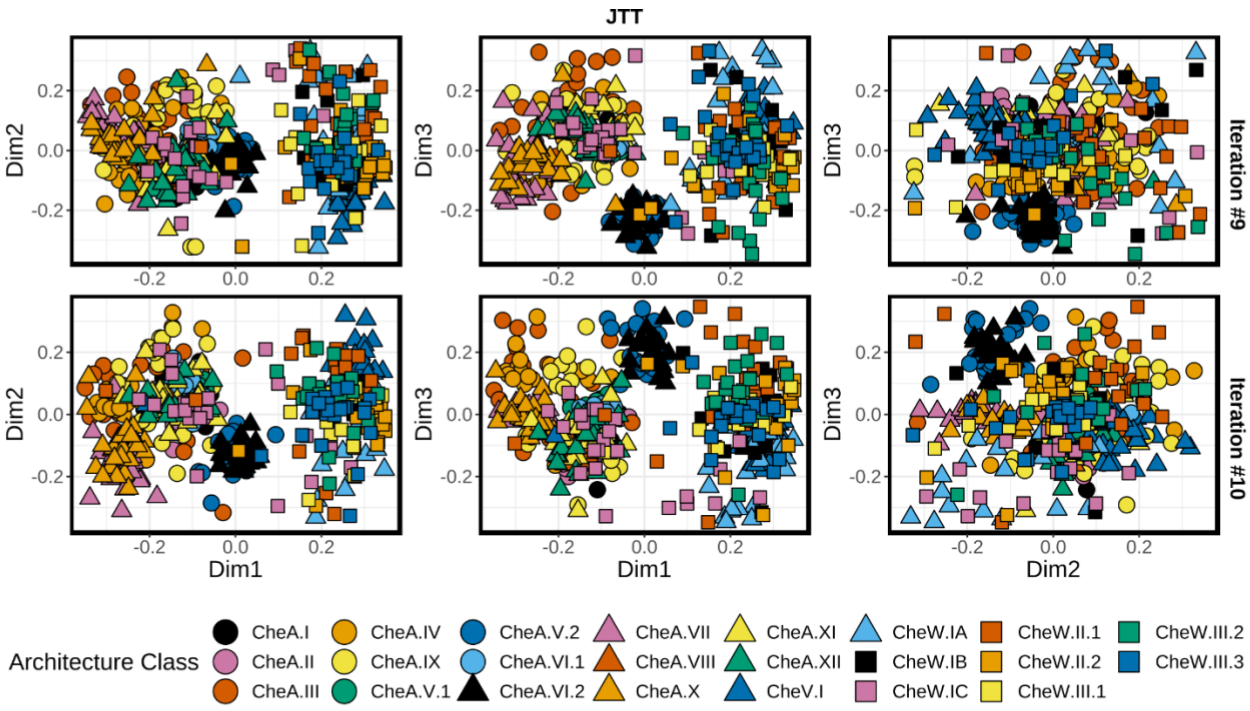

**FIGURE S20. BLOSUM30-derived nMDS solutions – Iterations #1-4**

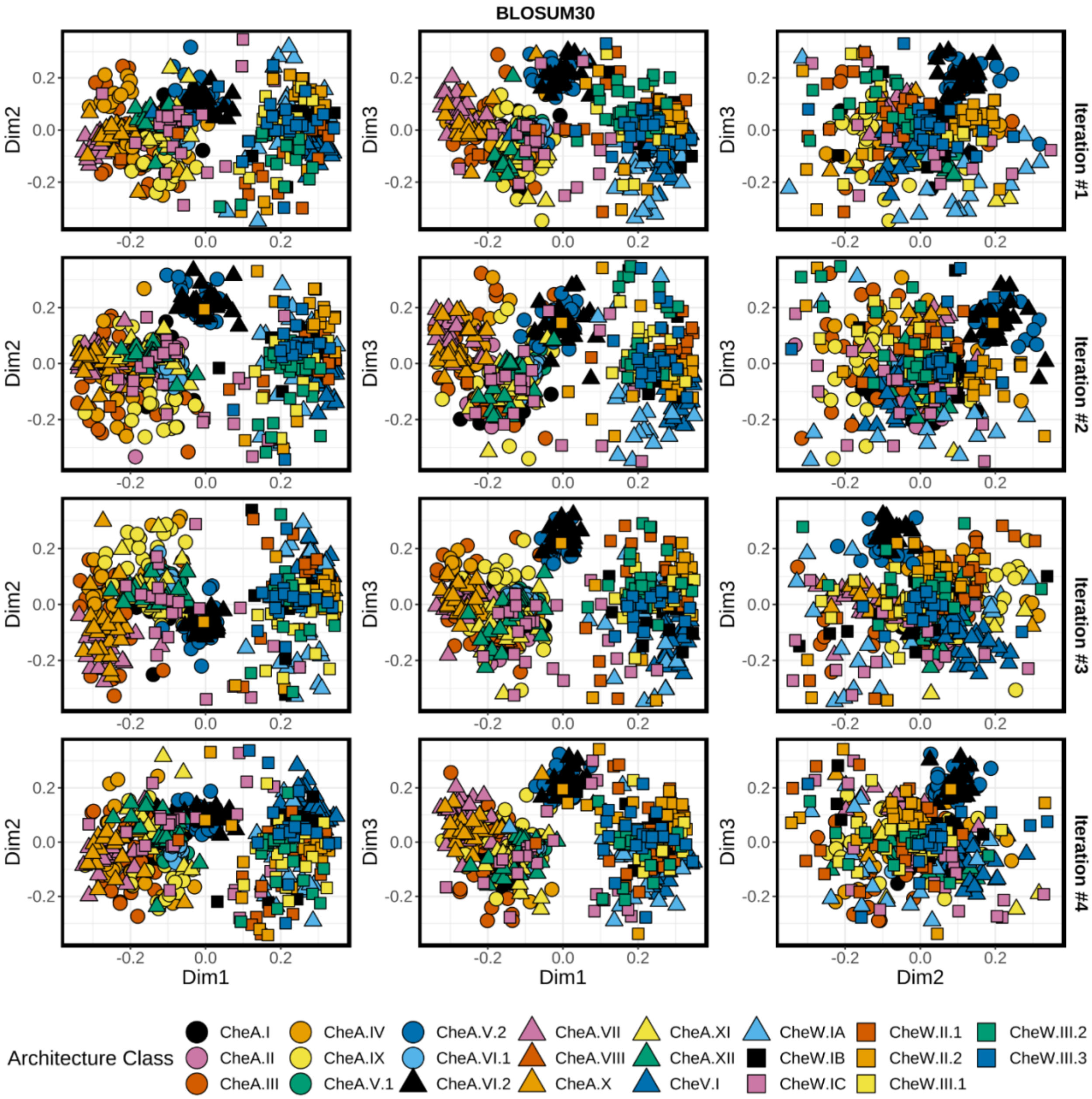

**FIGURE S21. BLOSUM30-derived nMDS solutions – Iterations #5-8**

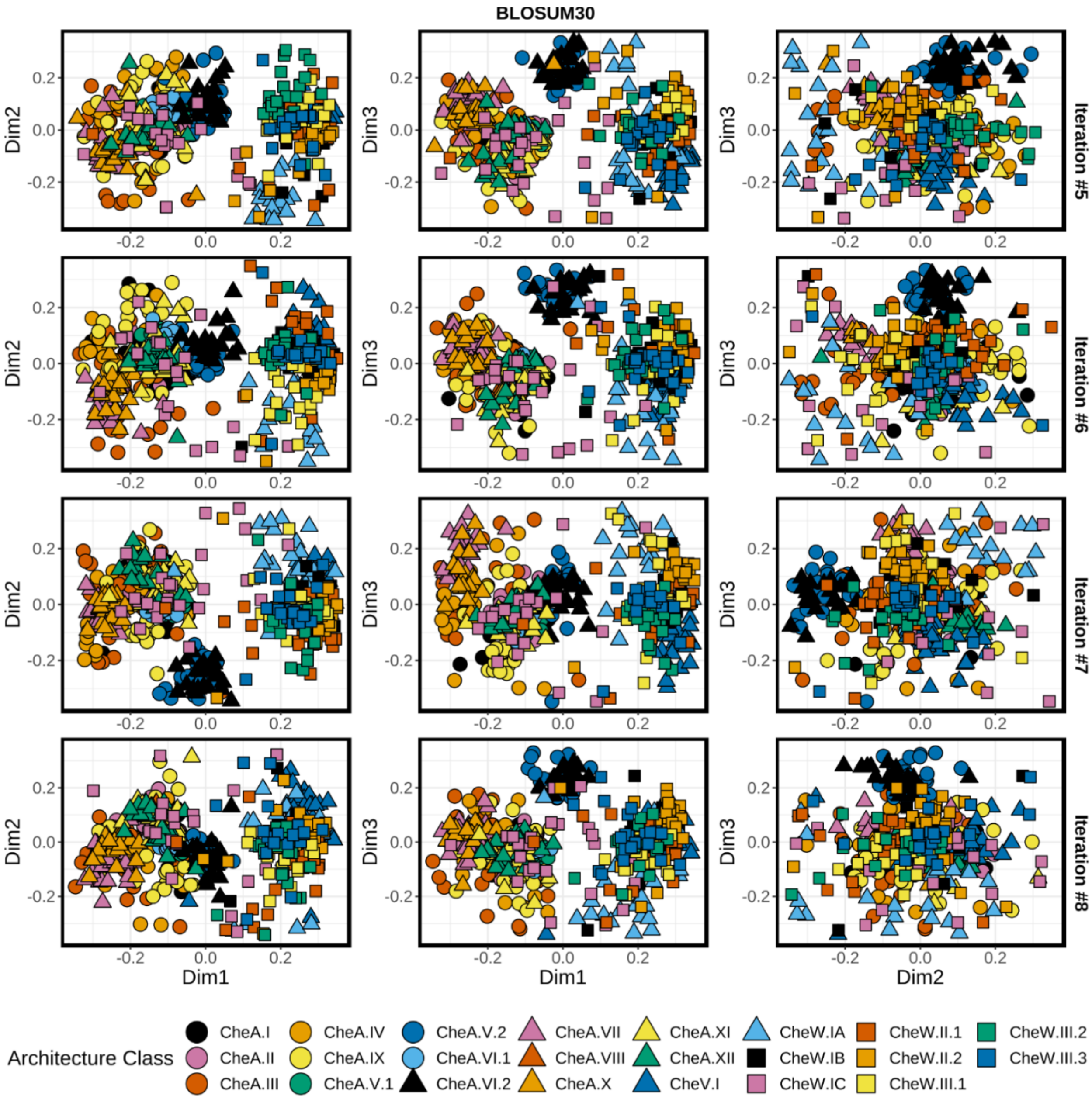

**FIGURE S22. BLOSUM30-derived nMDS solutions – Iterations #9-10**

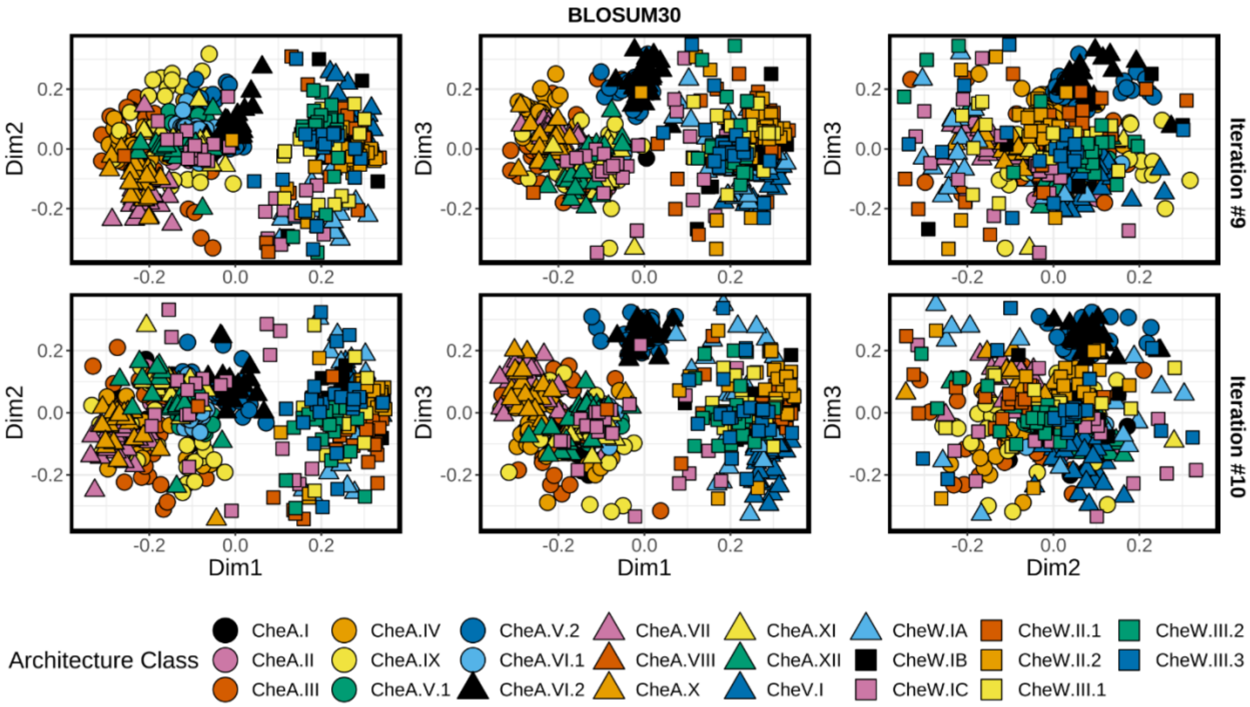

**FIGURE S23. PAM250-derived nMDS solutions – Iterations #1-4**

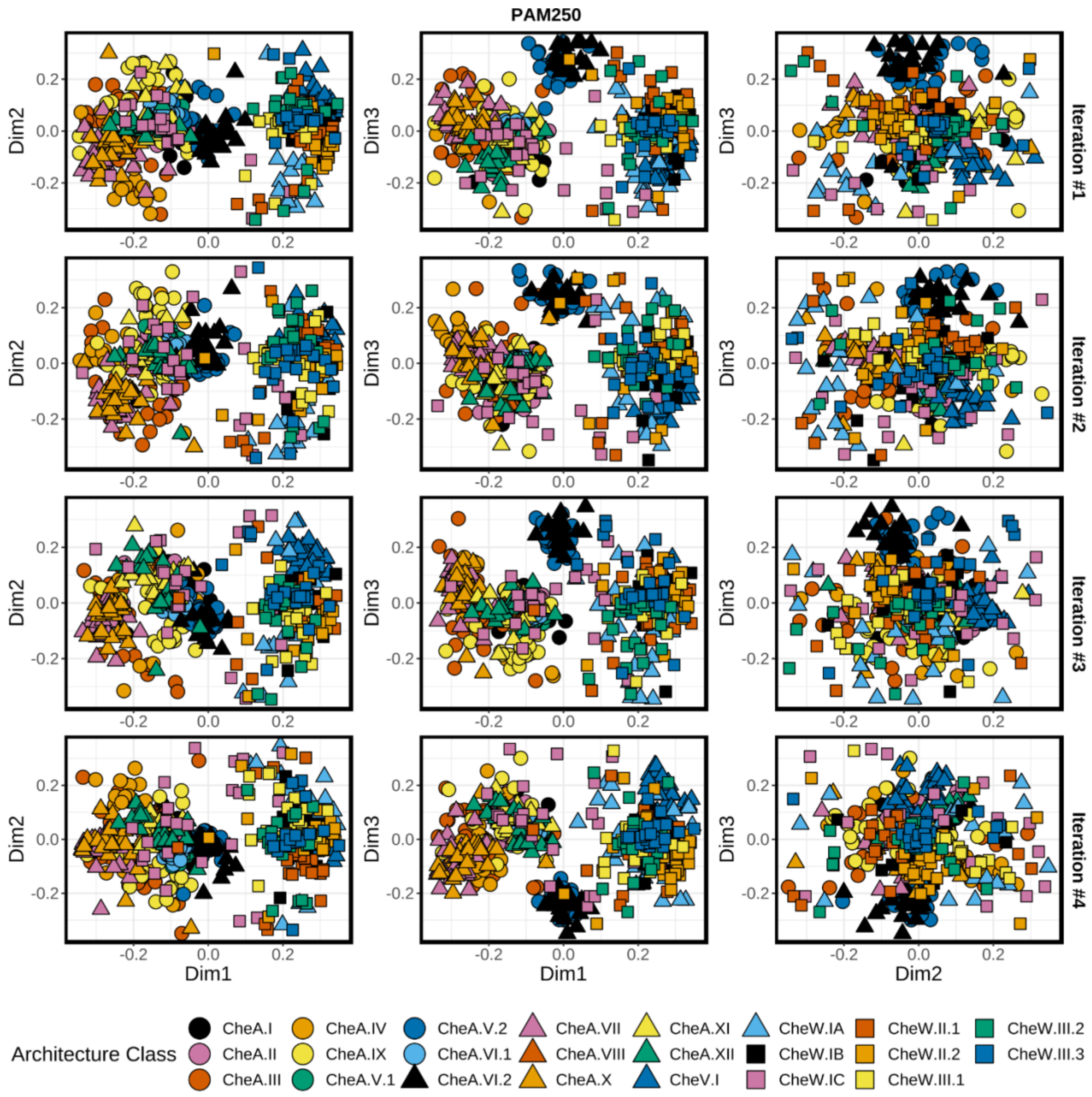

**FIGURE S24. PAM250-derived nMDS solutions – Iterations #5-8**

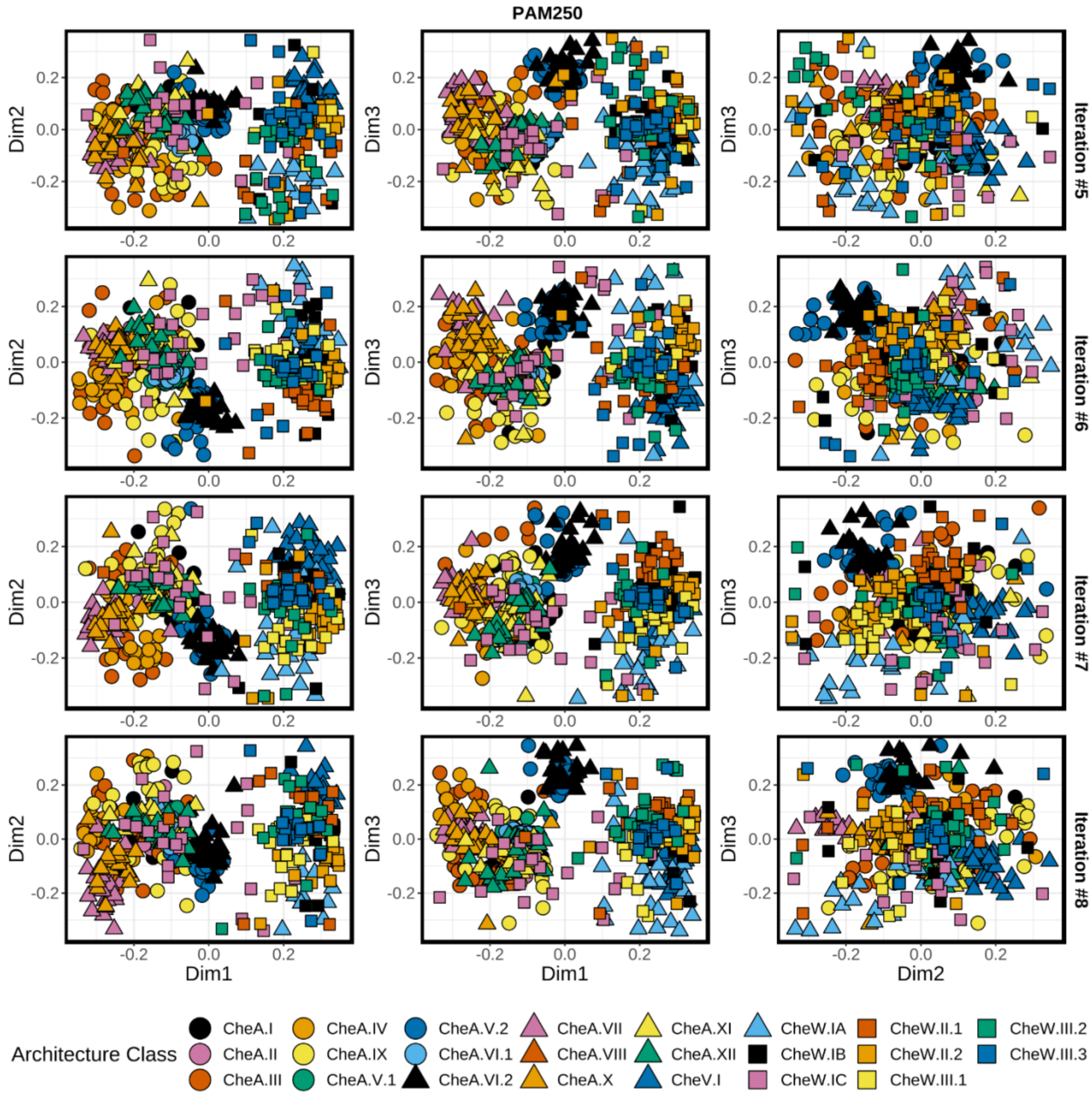

**FIGURE S25. PAM250-derived nMDS solutions – Iterations #9-10**

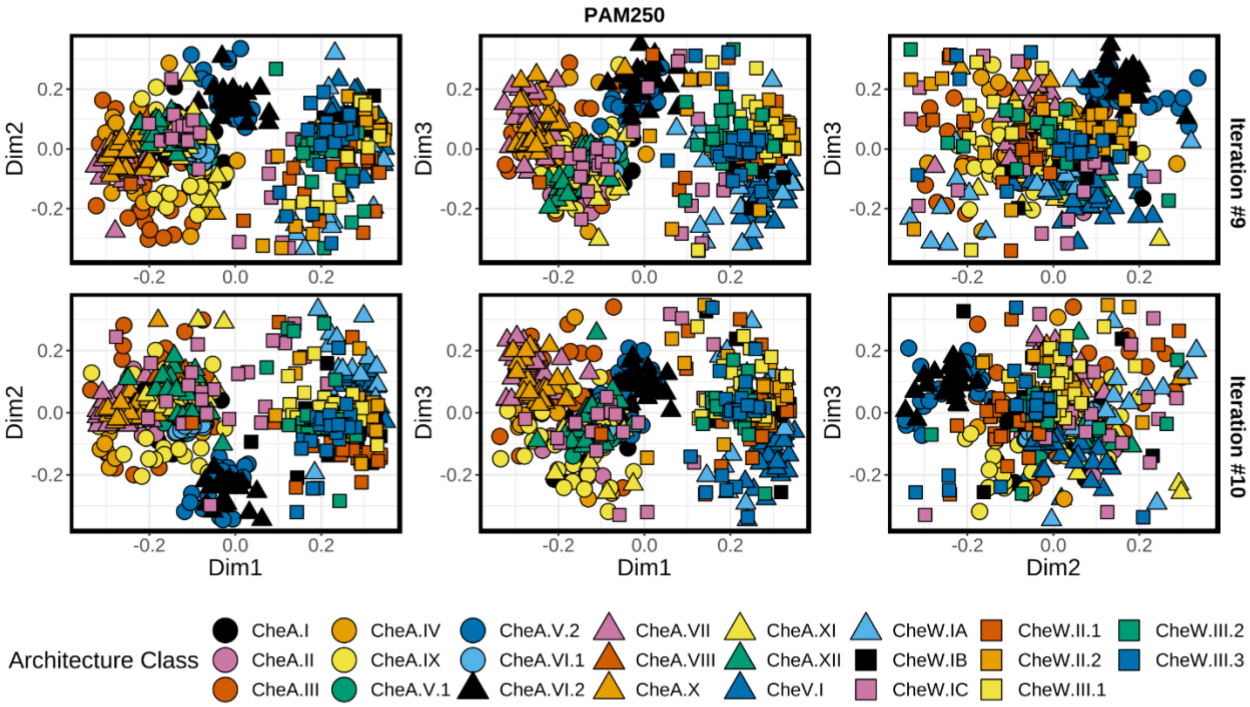

**FIGURE S26. Traditional sequence logo representation of the major CheW-like domain *Types* shown in Figure 4.** Rows correspond to domain *Types*. Sequences were aligned to the CheW-like domain HMM (Pfam ID PF01584) and separated by *Type*. Characters were colored based on amino acid type (default settings). Overall stack heights were scaled by information content (bits, scales not shown to simplify visualization) calculated at each position within the alignment, relating to conservation

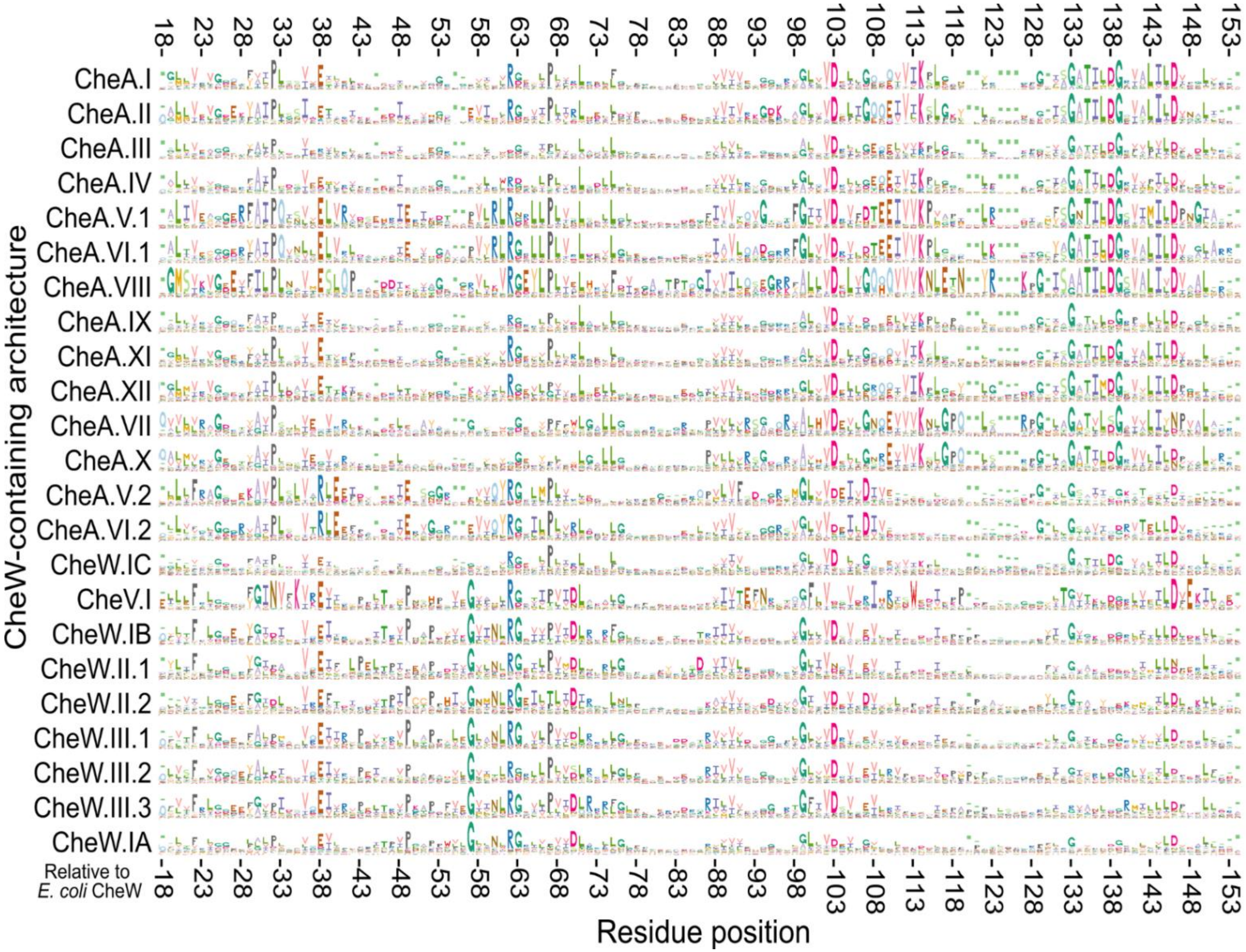
